# In vivo programming of stem-like CAR T cells by lymphatic-selective lipid nanoparticle enables durable anti-tumour efficacy in orthotopic model

**DOI:** 10.64898/2026.05.08.723693

**Authors:** Yishan Ma, Jingyi Chen, Xiaonan Huang, Jingsheng Cai, Guanghui Ma, Mantang Qiu, Yufei Xia

**Author notes:** These authors contributed equally to this work (Yishan Ma, Jingyi Chen, Xiaonan Huang).

## Abstract

Chimeric antigen receptor (CAR) T-cell therapy has shown remarkable efficacy in hematological malignancies, yet its efficacy in solid tumours remains limited by poor persistence and progressive exhaustion within the tumour microenvironment. These barriers may be particularly pronounced in emerging *in vivo* CAR-T therapies, in which transient transgene expression and insufficient control over T-cell differentiation restrict the generation of durable antitumour immunity. Here, we report a primary lymphoid tissue-targeting lipid nanoparticle (pLNP), that directs in vivo CAR-T programming to the thymus and lymphoid tissues, thereby increasing the proportion of stem-like CAR-T cells and promoting durable, exhaustion-resistant antitumour responses. After antibody conjugation, pLNP enabled in vivo CAR expression in developing T cells, generating CAR-T cells enriched in naïve and stem cell-like memory phenotypes with prolonged persistence. To reinforce this, we co-administered interleukin-7 (IL-7) mRNA, which increased stem-like CAR-T populations, favoured progenitor exhausted T (Tpex) cells over terminally exhausted states, and enhanced cytotoxic function without overt inflammatory amplification. This stemness-promoting strategy also improved responsiveness to immune checkpoint blockade, producing synergistic antitumour effects with anti-PD-1 therapy, reducing LNP dose requirements, and inducing durable tumour regression with prolonged survival in both subcutaneous and orthotopic DLL3-positive small-cell lung cancer models. Similar enhancement of in vivo CAR-T efficacy was also observed in aged mice with thymic involution. Together, these findings illustrated that primary lymphoid tissue-directed *in vivo* CAR-T programming is a potential strategy to overcome insufficient persistence and progressive exhaustion in solid tumours.

## Introduction

Chimeric antigen receptor T-cell (CAR T) therapy has achieved transformative clinical success in hematological malignancies, with six FDA-approved products for diseases, including acute lymphoblastic leukemia and B-cell lymphomas^1–3^. Yet, current CAR T manufacturing relies on extensive *ex vivo* T-cell activation, expansion and genetic modification, processes that can erode T-cell stemness and drive terminal differentiation^4–6^. *In vivo* programming of immune cells offers an alternative route^7–12^, and antibody-conjugated mRNA lipid nanoparticles (LNPs) have emerged as a representative platform that can simplify manufacturing, reduce time-to-treatment, and enable repeated dosing without complex cell-processing infrastructure^9,14–17^. However, durable therapeutic efficacy remains difficult to achieve^18^, owing not only to the transient nature of mRNA expression, but also to the lack of deliberate control over T-cell differentiation toward long-lived antitumour states^19–24^. These limitations are further amplified in solid tumours by an immunosuppressive tumour microenvironment (TME) that restricts T-cell infiltration, impairs intratumoural function, and drives progressive exhaustion under chronic antigen exposure^25–29^. Thus, for next-generation *in vivo* CAR-T treatment against solid tumours, durable benefit will likely require not only efficient gene delivery but also sustained T-cell persistence and resistance to terminal exhaustion^29^.

Increasing evidence indicates that “stem-like” T-cell states play a central role in sustaining antitumour immunity during chronic antigen exposure. Within the exhausted T-cell continuum, progenitor exhausted T (Tpex) cells represent a less differentiated subset with self-renewal capacity, sustained proliferative potential, and responsiveness to cytokine cues, and have recently been discovered as the major T-cell population mediating responses to and immune checkpoint signaling, such as PD-1/PD-L1 antibodies in TME^30–34^. By contrast, terminally exhausted T (Tex) cells display limited proliferative potential and diminished functional rescue. More broadly, durable CAR-T responses appear to rely on stem-like T-cell populations, ranging from stem-like memory T cells (Tscm), which are associated with early persistence, to Tpex cells that propagate and replenish effector cells under chronic antigen exposure in TME^31,35–39^. These observations suggest that the central challenge is not simply to generate CAR-T cells *in vivo*, but to bias their developmental trajectory toward stem-like states that can persist, self-renew, and withstand exhaustion pressure in solid tumours.

We therefore hypothesized that *in vivo* engineering of immature, developmental T cells may intrinsically favour stem-like CAR T cells with greater durability. The thymus, as the central organ for T-cell development, selection and tolerance formation, provides a highly regulated microenvironment for T-cell lineage commitment, activation and differentiation^41–44^. This primary immune niche suggests a unique opportunity for *in vivo* CAR-T strategy: thymus- or primary-lymphatic-tissue-targeted *in vivo* programming may favour the generation of CAR-T cells with improved stemness and longevity, compared with approaches that predominantly engineer peripherally activated T cells. In parallel, interleukin-7 (IL-7), a key cytokine for T-cell development, survival, and homeostasis, supporting the maintenance of naïve and stem-like T-cell populations^45–50^, has been associated with enhanced T-cell persistence, preserved proliferative capacity, and restrained terminal exhaustion^37,51–52^. Together, these considerations led us to test whether combining primary lymphoid tissue-directed CAR engineering armoured with IL-7 supplementation could establish a more stem-like and durable CAR-T state, with the potential to improve antitumour immunity and responsiveness to immune checkpoint blockade.

In this study, we engineered a primary lymphoid tissue-targeting lipid nanoparticle (pLNP) platform for *in vivo* CAR-T programming. We further co-administered IL-7 mRNA-loaded LNPs to reinforce stem-like and functional CAR-T states, thereby enhancing persistence and improving responsiveness to immune checkpoint blockade in both subcutaneous and orthotopic lung tumour models. Similar enhancement of antitumour activity was also observed in aged mice with thymic involution (data are not shown). By integrating spatially directed CAR engineering with cytokine support, this strategy addresses two central bottlenecks of current *in vivo* CAR-T approaches for solid tumours – limited long-term persistence and progressive exhaustion within the tumour microenvironment -- and thereby improves sustained antitumour efficacy.

## Result

### LNP screening for mRNA delivery to T cells in lymphatic tissues

To identify LNP formulations with lymphoid organ tropism, we constructed and screened a panel of LNPs. Two benchmark lipids, ALC0315 (BNT162b2 vaccine), LP-01 (US11420933B2), together with two lipids previously developed in our laboratory, OT13^19^ and GT7 (WO2025256311A1), were selected as the ionizable lipid component. Using these four ionizable lipids together with two helper phospholipids (DSPC and DOPE), two PEG-lipid variants (DSPE-PEG2000 and DMG-PEG2000), and three classical four-component LNP compositions (30:46.5:16:2.5, 40:48.5:10:1.5, and 45:44:9:2), we assembled a 48-member LNP library for in vivo screening (**Fig. 1a and Supplementary Fig. 1a**).

**Fig. 1.**
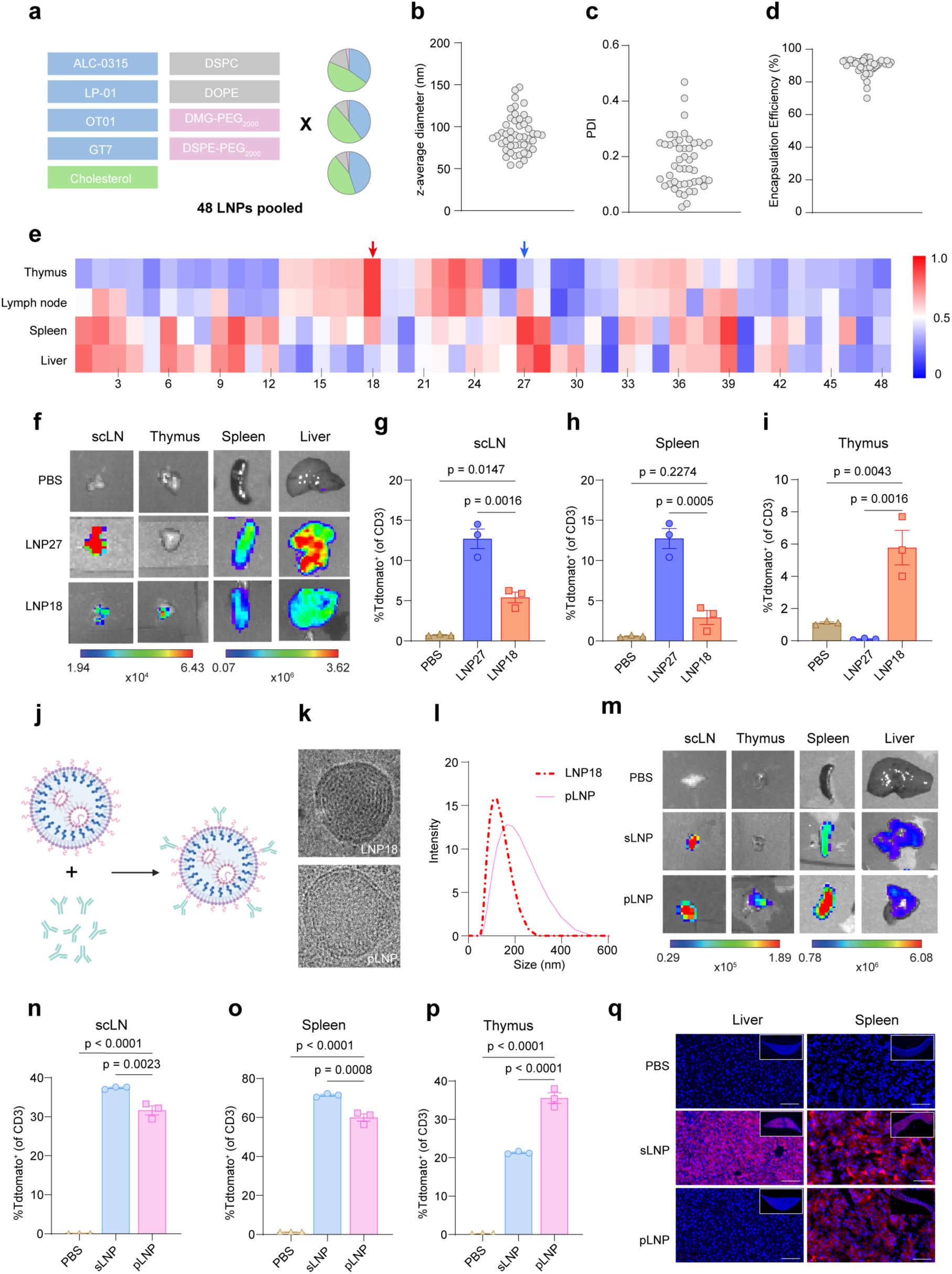
LNP formulation screening for in vivo T-cell transfection in primary and secondary lymphoid tissues. **a**, Schematic illustration of the LNP design with different ionizable lipids, helper lipid, PEGylated lipids and component ratios. **b-d**, Characterization of 48 distinct LNPs. Each LNP was analyzed for hydrodynamic diameter (**b**), Polydispersity Index (PDI) (**c**), mRNA encapsulation efficiency (**d**). **e**, In vivo bioluminescence heatmap of luciferase expression across thymus, lymph node, spleen and liver. **f**, Representative IVIS images of dissected organs following administration of PBS (negative control), LNP27 (spleen-tropic LNP), LNP18 (thymus-tropic LNP). **g–i**, Cre-mediated tdTomato reporter activation in Ai9 mice 5 days after injection of LNP27 or LNP18 encoding Cre mRNA, showing differential T-cell transfection efficiency in lymph node (**g**), spleen (**h**), and thymus (**i**). **j**, Schematic illustrating antibody conjugation to LNPs for targeted in vivo CAR T-cell generation. **k**, Cryo-TEM imaging of LNP18 before and after antibody conjugation. **l**, Hydrodynamic diameter analysis of LNP18 before and after antibody conjugation. **m**, Representative IVIS images of organs following i.v. injection of sLNP or pLNP at dose of 0.2 mg kg^−1^. **n–p**, Quantification of TdTomato⁺ CD3⁺ T cells in lymph node (**n**), spleen (**o**), and thymus (**p**) in Ai9 mice 5 days after injection of Cre-encoding sLNP or pLNP, demonstrating preferential thymic targeting by pLNP and enhanced T-cell transfection in lymphatic tissues. **q**, Cryo section of Liver and spleen after i.v. injection of sLNP and pLNP expressing tdTomato. Data are presented as mean ± s.e.m. Statistical significance was determined by one-way ANOVA with multiple comparisons. p values are indicated in the panels.

The resulting LNPs exhibited generally comparable physicochemical characteristics, with z-average diameters mainly in the ∼60–150 nm range, PDI values distributed broadly but generally below 0.3, and encapsulation efficiencies mostly around 80–95% (**Fig. 1b–d**). All 48 LNPs were then pooled and intravenously administered to wild-type mice to evaluate their ability to deliver luciferase mRNA *in vivo*. Six hours after administration, bioluminescence imaging was performed in lymphatic tissues, including spleen, superficial cervical lymph node (as termed scLNs), and thymus, as well as liver. The averaged flux from respective tissue was quantified and visualised using heatmaps (**Fig. 1e**). Among the screened formulations, LNP18 showed preferential luciferase expression in lymphoid organs, particularly the thymus, whereas LNP27 showed predominant expression in the spleen, lymph node, and liver, with minimal signal detected in the thymus (**Fig. 1e, f**).

To define the cellular tropism of the selected formulations in lymphatic organs, we next evaluated Cre mRNA delivery in genetically engineered Ai9 tdTomato reporter mice carrying a LoxP-flanked stop cassette, in which Cre expression excises the stop cassette and enables fluorescent tdTomato expression in transfected cells. LNP27 showed higher tdTomato expression in CD3⁺ T cells from the lymph node and spleen, whereas LNP18 preferentially transfected thymic T cells (**Fig. 1g–i).**

We next employed antibody conjugation to facilitate cell-selective delivery for in vivo CAR-T generation. We chose the anti-CD5 antibody because CD5 is naturally expressed on T cells and has minimal impact on T-cell homeostasis^8^. Accordingly, we engineered two aCD5-conjugated formulations derived from LNP27 and LNP18, termed sLNP (secondary lymphoid organs targeting LNP) and pLNP (primary lymphoid organs targeting LNP), respectively (**Fig. 1j**) ^8,14–16^. Compared with LNP27 and LNP18, sLNP and pLNP formulations showed an approximately 50 nm increase in hydrodynamic diameter after conjugation, without obvious structural changes under cryo-TEM evaluation (**Fig. 1k-l, Supplementary Fig. 1b-c**). Antibody conjugation markedly shifted the biodistribution of both formulations toward lymphoid tissues and away from the liver relative to their corresponding unconjugated formulation (**Fig. 1m and Supplementary Fig. 1d**).

In Ai9 reporter mice, sLNP mediated higher tdTomato expression in T cells in the secondary lymphoid organs (scLN and spleen), whereas pLNP more effectively transfected thymic T cells **(Fig. 1n-p)**. Moreover, compared with LNP18, pLNP showed reduced transfection of non-T-cell populations, including B cells, macrophages, dendritic cells, and myeloid cells, indicating that antibody conjugation improved T-cell selectivity (**Supplementary Fig. 1e**). In addition, pLNP-mediated transgene expression remained negligible in the liver, indicating limited off-target tissue distribution (**Fig**. **1q**). Together, these results demonstrated two antibody-conjugated LNP platforms with distinct lymphoid-organ tropism. sLNP targeted T cells in the secondary lymphoid tissues, including the lymph node and spleen, whereas pLNP enables delivery to thymic T cells in the primary lymphoid organ while also retaining access to secondary lymphoid tissues.

### Enhancing T-cell stemness to promote durable *in vivo* CAR-T dynamics and antitumour efficacy

We next used sLNP and pLNP to encapsulate a circular RNA encoding a DLL3 (Delta-like ligand 3) CAR (sLNP^CAR^ and pLNP^CAR^). DLL3 is an atypical Notch ligand aberrantly overexpressed on the surface of small-cell lung cancer (SCLC). The construct and the corresponding circular RNA have previously been designed and validated^20^ (**Fig. 2a**).

**Fig. 2.**
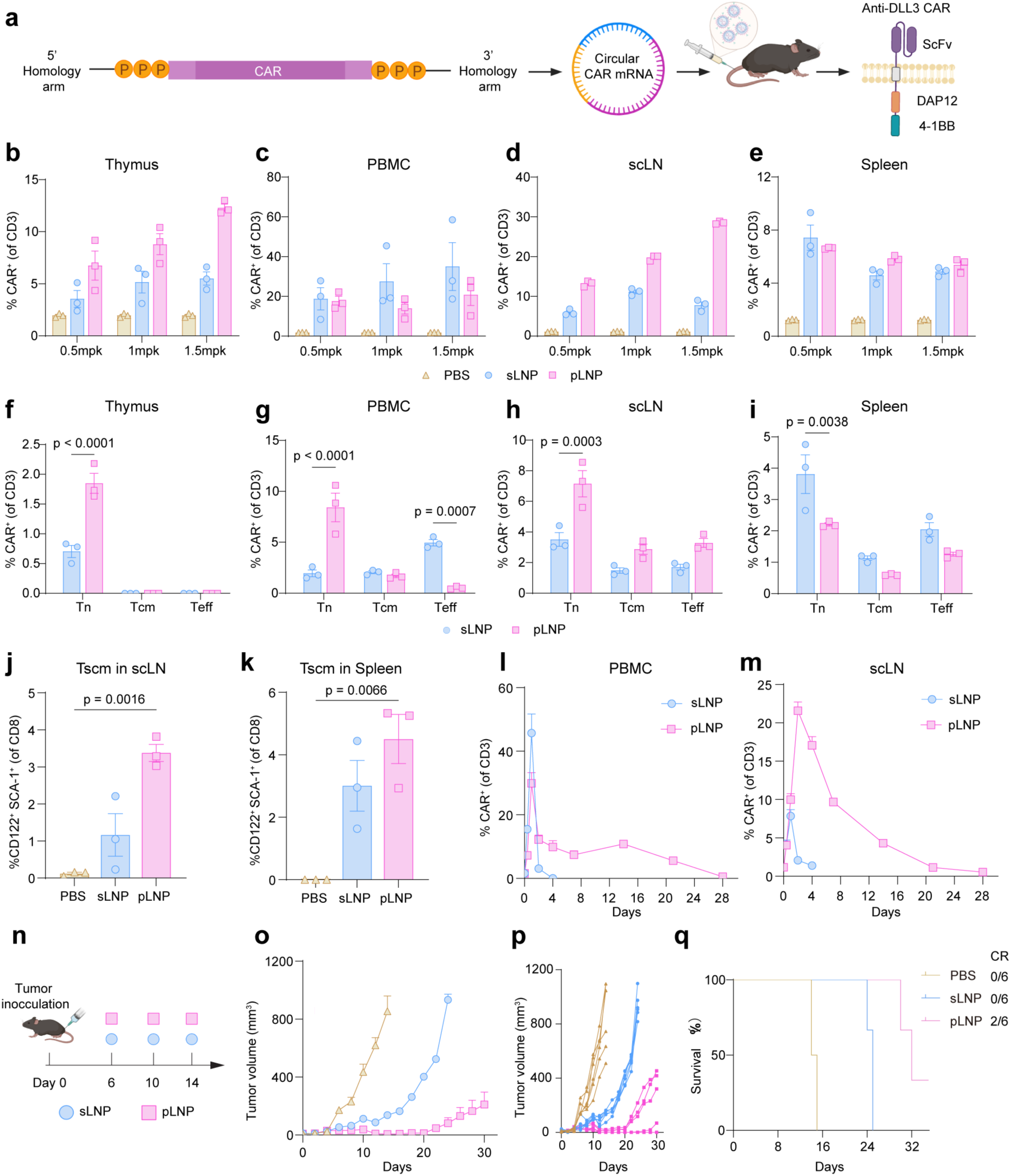
pLNP mediated *in vivo* CAR-T promotes a naïve-biased differentiation associated with improved antitumour efficacy. **a,** Schematic illustrating the generation of circular CAR mRNA encoding an anti-DLL3 chimeric antigen receptor and its delivery in vivo using antibody-conjugated lipid nanoparticles (LNPs). **b–e**, Quantification of CAR expression among CD3⁺ T cells 48h after i.v. administration of sLNP or pLNP at doses of 0.5, 1, or 1.5 mg kg⁻¹. CAR⁺ T-cell frequencies were measured in the thymus (**b**), peripheral blood mononuclear cells (PBMCs) (**c**), lymph nodes (**d**), and spleen (**e**). **f–i**, Phenotypic profiling of CAR⁺ T cells 48 h after LNP administration. The relative proportions of naïve T cells (Tn), central memory T cells (Tcm), and effector T cells (Teff) subsets were analyzed among CAR⁺ CD3⁺ T cells in thymus (**f**), PBMCs (**g**), lymph nodes (**h**), and spleen (**i**). pLNP delivery preferentially enriched T naïve cells compared with sLNP. **j-k**, Frequency of stem cell–like memory T cells (Tscm; CD62L⁺CD122⁺Sca1⁺) among CD8⁺ T cells in lymph nodes (**j**) and spleen (**k**) following LNP treatment. Gating strategy refers to **Supplementary** Fig. 6. **l,m**, Longitudinal analysis of CAR⁺ T cells in PBMCs (**l**) and lymph nodes (**m**), at indicated time points, demonstrating sustained CAR expression following pLNP-mediated delivery compared with sLNP. **n**, Experimental design of the DLL3^+^ subcutaneous tumour model following administration of sLNP or pLNP. **o-p**, Tumour growth curve measured every 2 days (**o**) and individual tumour growth curve treated with PBS, sLNP, or pLNP (**p**). **q**, Probability of survival showing improved survival following pLNP treatment, including complete responses (CR), 2 out of 6 observed. Statistical significance was determined by one-way ANOVA with multiple comparisons. p values are indicated in the panels.

Further, we asked whether thymus-accessible CAR delivery could influence the differentiation state of in vivo-generated CAR-T cells. To address this, we analysed the abundance and phenotype of CAR⁺ T cells in the thymus, scLN, spleen, and peripheral blood after intravenous administration of sLNP^CAR^ or pLNP^CAR^ at the highest expression dose of 1.5 mg kg^−1^ (**Fig. 2b-e**), with tissues collected at 48 h. Across these anatomical sites, phenotypic profiling revealed distinct differentiation and circulation states of the generated CAR-T cells (**Fig. 2f–i**). Notably, pLNP^CAR^ preferentially enriched CAR⁺ naïve T cells (Tn) in the thymus, PBMCs, and lymph nodes, whereas sLNP^CAR^ showed a relatively greater representation of CAR⁺ effector T cells in PBMCs and spleen.

Because the thymic niche is closely linked to early T-cell developmental programming, we hypothesized that directing CAR expressions towards these sites, rather than the periphery, might more efficiently skew their differentiation toward a T stem-like memory cells (Tscm) phenotype, which has been associated with sustained CAR-T persistence. As shown in **Fig. 2j, k**, pLNP^CAR^ yielded a higher proportion of CAR⁺ Tscm cells in both lymph nodes and spleen, compared to sLNP^CAR^. Additionally, pLNP^CAR^ induced comparable CAR expression in the T cells in the peripheral blood and spleen with sLNP^CAR^ but produced substantially higher CAR^+^ T-cell frequencies in the thymus and lymph nodes in a clear dose-dependent manner, indicating enhanced CAR expression in both primary and secondary lymphoid tissues (**Fig. 2b–e**).

Notably, longitudinal analysis further showed that pLNP sustained CAR expression for up to 28 days, particularly in lymph nodes (**Fig. 2l, m**). At 48 h, the frequency of CAR⁺ T cells in lymph nodes were approximately 10-fold higher in the pLNP group than in the sLNP group, whereas sLNP-mediated expression was largely confined to PBMCs and declined within 4 days.

We then evaluated whether this stemness-associated shift translated into improved anti-tumour activity in vivo. In an established subcutaneous LLC-DLL3-Luc tumour model (**Fig. 2n**). pLNP^CAR^ induced greater tumour regression than sLNP^CAR^ (**Fig. 2o, p**), achieving complete responses (CR) in 33% (2 out of 6) of the treated mice and significantly extending median survival compared to the sLNP^CAR^ cohort (**Fig. 2q**).

We further validated these findings in a more stringent orthotopic tumour model, where pLNP^CAR^ similarly outperformed sLNP^CAR^ in controlling tumour burden and prolonging overall survival (**Supplementary Fig. 2a-c**). In addition, both sLNP^CAR^ and pLNP^CAR^ upregulated intratumoural TNF-α and IFN-γ compared with PBS (**Supplementary Fig. 2d, e**). However, this robust initial effector function was accompanied by a marked accumulation of immunosuppressive regulatory T cells (Tregs) within the tumour microenvironment (**Supplementary Fig. 2f**). Such a suppressive response under continuous antigen exposure may drive CAR-T cell exhaustion and compromise long-term therapeutic efficacy. But pLNP^CAR^ induced significantly lower intratumoural Treg than sLNP^CAR^ (p = 0.0052), indicating that a stemness-favouring CAR-T state may better resist exhaustion-associated dysfunction. Additionally, H&E staining confirmed the safety of pLNP (**Supplementary Fig. 2g)**.

Collectively, these findings suggest that extending CAR delivery beyond secondary lymphoid tissues to the thymus can reshape in vivo CAR-T dynamics toward naïve- and stem-like-dominated populations. This shift promotes CAR-T accumulation and persistence in major lymphoid organs and is associated with reduced exhaustion-prone suppressive pressure and improved antitumour efficacy.

### IL-7 co-delivery for boosted durability and stemness of CAR-T cells in vivo

Given the enhanced lymphatic targeting and prolonged CAR expression mediated by pLNP, we next investigated whether this platform could be further leveraged to support CAR-T persistence through in situ cytokine supplementation (**Fig. 3a**). LNP18 encapsulating luciferase mRNA was subcutaneously injected into C57BL/6J mice, and bioluminescence imaging then demonstrated preferential expression within the lymph node with limited liver exposure (**Fig. 3b**). This lymph node-oriented expression suggested that LNP18 could serve as a platform for local cytokine support of CAR-T cells.

**Fig. 3.**
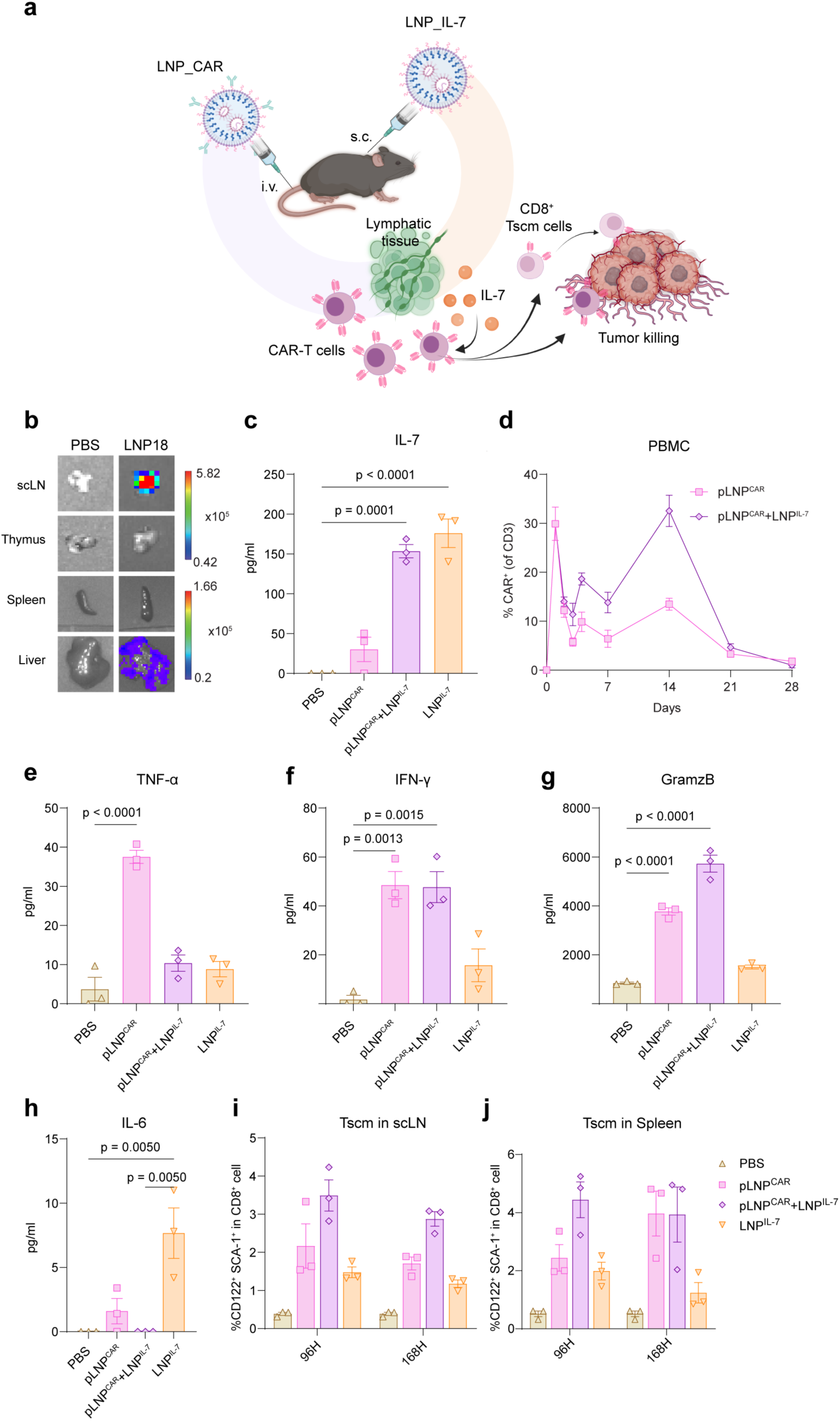
Combination therapy with LNP^IL-7^ enhances stem-like CAR T cell programming and promotes cytotoxicity. **a**, Schematic illustration of the combination strategy image created with BioRender. pLNP^CAR^ were administered intravenously to induce in vivo CAR expression, followed by subcutaneous injection of IL-7 mRNA encapsulated LNPs to establish a lymphatic cytokine niche supporting CAR T-cell persistence and function. **b**, Ex vivo bioluminescence imaging of major organs after subcutaneous delivery of luciferase mRNA encapsulated in LNP18, showing preferential signal localization in lymphatic tissues with minimal hepatic exposure. H, heart; Lu, lung; K, kidney; Li, liver; S, spleen; Ln, lymph node; Th, thymus. **c**, IL-7 secretion in LN was confirmed by Elisa with co-administration of pLNP^CAR^ and LNP^IL-7^ or LNP^IL-7^ alone groups**. d**, Longitudinal quantification of CAR⁺ T cells in peripheral blood mononuclear cells (PBMCs) following pLNP^CAR^ alone or in combination with LNP^IL-7^, demonstrating enhanced and sustained CAR expression in the combination group. **e-h**, cytokine levels were evaluated from mouse lymph node homogenates by ELISA. TNF-α (**e**), IFN-γ (**f**), granzyme B (**g**), and IL-6 (**h**). **i-j**, Frequency of stem cell– l ike memory T (TSCM) cells (CD44⁻CD62L⁺Sca-1⁺CD122⁺ within CD8⁺ T cells) in lymph nodes (**i**) and spleen (**j**) at 96 h and 168 h post-treatment. IL-7 co-administration induced a transient expansion of TSCM cells that remained elevated relative to controls. Data are presented as mean ± s.e.m. Statistical significance was determined by one-way ANOVA with multiple comparisons. p values are indicated in the panels.

We therefore screened several candidate cytokines ex vivo, including IL-2, IL-4, IL-7 and IL-12. *Ex vivo* comparison showed that IL-7 provided the most favourable overall balance among T-cell viability, proliferation, and exhaustion-associated phenotypes, and was therefore selected for subsequent studies (**Supplementary Fig. 3a–c**).

We then assessed whether IL-7–mediated conditioning could reinforce a stem-like CAR T cell state and enhance persistence^45–49,52^. IL-7 mRNA was encapsulated in LNP18 and administered subcutaneously 24 h following pLNP-mediated *in vivo* CAR engineering. ELISA analysis of scLN confirmed robust IL-7 production following coadministration of LNP^IL-7^ (**Fig. 3c).** Longitudinal analysis revealed combinatorial treatment of pLNP^CAR^ and LNP^IL-7^ significantly enhanced CAR expression in peripheral blood, reaching a peak at day 14 and remaining detectable through day 28 (**Fig. 3d**), while not further increasing CAR expression within LN (**Supplementary Fig. 3d**).

Compared with pLNP^CAR^ alone, co-administration of LNP^IL-7^ increased TNF-α, IFN-γ, and granzyme B levels (**Fig. 3e-g**), indicating enhanced cytotoxic effector function of CAR-T cells, without evidence of excessive inflammatory activation, as IL-6 remained low (**Fig. 3h**). Flow cytometry further showed a transient increase in IL-2 production by CD8⁺ CAR-T cells, peaking at 96 h and returning toward baseline by 168 h (**Supplementary Fig. 3e, f**).

In addition, flow cytometric analysis revealed that, compared with pLNP^CAR^ alone, co-administration of pLNP^CAR^ and LNP^IL7^ significantly increased the frequency of CD8⁺ Tscm cells in the lymph nodes at both 96 h and 168 h, with a similar trend observed in the spleen (**Fig. 3i-j**). Together, these findings indicate that co-treatment with LNP^IL7^ enhances pLNP^CAR-^mediated CAR-T expression, durability, and cytotoxic activity *in vivo*, while also promoting a more stem-like CAR-T state that may contribute to improved persistence.

### Combination treatment of pLNP^CAR^ and LNP^IL-7^ exhibits a durable antitumour response and reshapes the tumour microenvironment in a DLL3 orthotopic tumour model

To further evaluate in vivo therapeutic efficacy, we next tested the co-administration of pLNP^CAR^ and LNP^IL-7^ in our established DLL3 SCLC orthotopic model. Mice were orthotopically injected with an aggressive, ex vivo passaged, LLC-DLL3 SCLC cell line expressing firefly luciferase (LLC-DLL3_luc). 14 days following tumour inoculation, mice received intravenous administration of pLNP^CAR^, followed 24 h later by subcutaneous administration of LNP^IL-7^ (**Fig. 4a**).

**Fig. 4.**
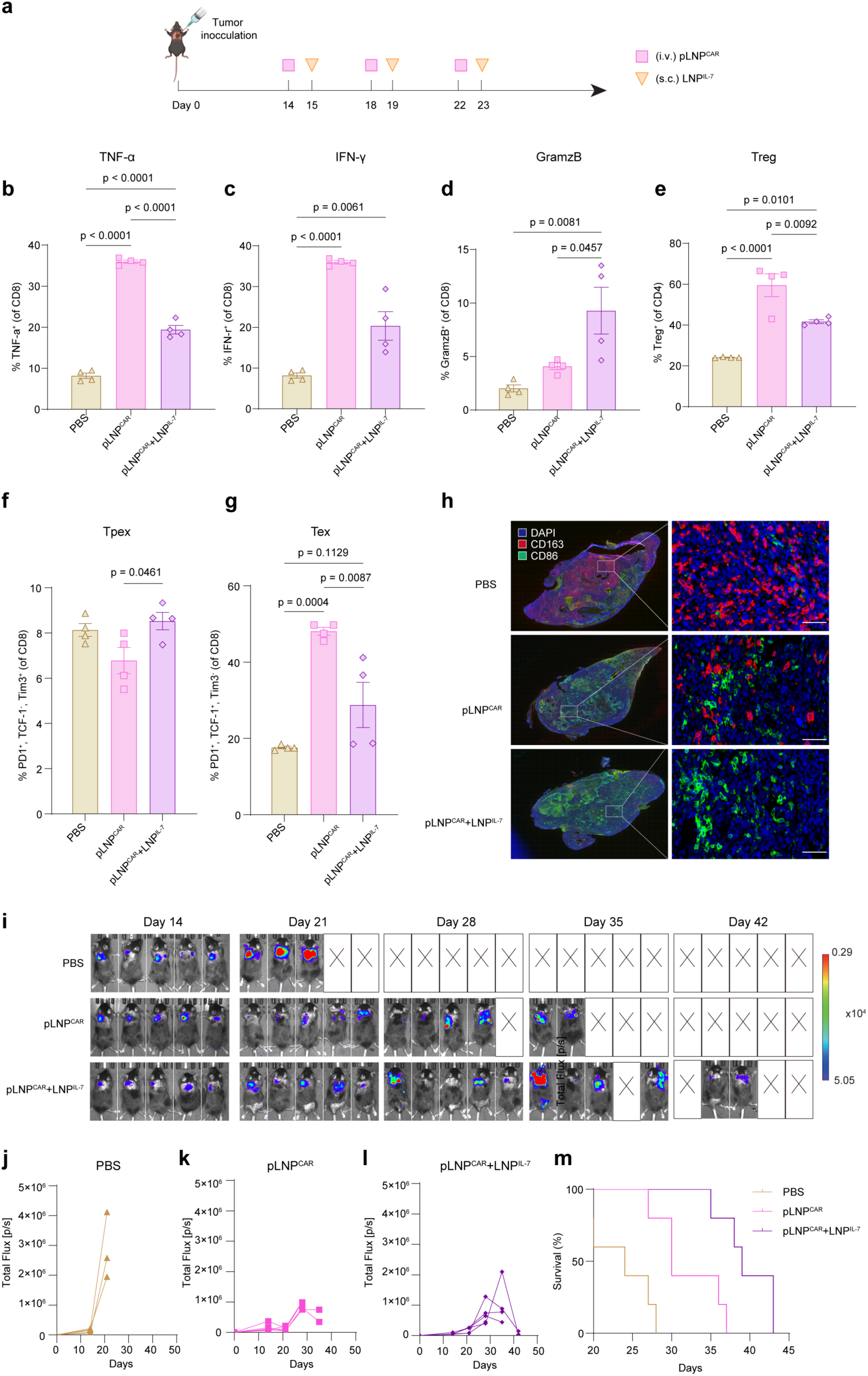
Combination therapy of pLNP^CAR^ and LNP^IL-7^ enhanced anti-tumour efficacy and remodeled tumour microenvironment in an orthotopic SCLC model. **a**, Experimental timeline. Mice bearing orthotopic DLL3⁺ SCLC tumours received intravenous pLNP^CAR^ followed by subcutaneous administration of LNP^IL-7^ at the indicated time points. **b-c**, Flow cytometric analysis of intratumoural CD8⁺ T cells. pLNP treatment increased inflammatory cytokine production (TNF-α (**b**) and IFN-γ (**c**)), whereas LNP^IL-7^ co-administration restrained excessive cytokine output while significantly enhancing granzyme B (**d**) expression, indicating a shift toward contact-dependent cytotoxicity. **e**, Frequency of regulatory T cells (Treg) within the tumour microenvironment. **f-g**, Quantification of progenitor exhausted (Tpex) (**f**) and terminally exhausted (Tex) CD8⁺ T-cell (**g**) subsets within tumours. Gating strategy refers to **Supplementary** Fig. 6. **h**, Immunofluorescence analysis of tumour sections stained for CD86 and CD163. Combination therapy increased CD86⁺ antigen-presenting cell infiltration and reduced CD163⁺ macrophage-associated signals. Scale bars, 50 µm. **i**, Representative bioluminescence imaging of tumour burden at days 14, 21, 28, 35, and 42 after tumour inoculation. pLNP^CAR^+LNP^IL-7^ resulted in sustained tumour regression compared with PBS and pLNP alone. **j-l**, Quantification of individual tumour burden bioluminescence signals, PBS (**j**), pLNP^CAR^ (**k**), pLNP^CAR^+LNP^IL-7^ (**l**). **m**, Survival rate demonstrating significantly prolonged survival in the combination group relative to controls. Statistical significance was determined by one-way ANOVA with multiple comparisons. p values are indicated in the panels.

To profile the tumour microenvironment (TME), we then analyzed tumour-infiltrating lymphocytes using FACS analysis. Within tumour-infiltrating CD8⁺ T cells, TNF-α and IFN-γ production was reduced in the co-administration group relative to pLNP^CAR^ alone (**Fig. 4b, c**), whereas granzyme B expression was significantly increased (**Fig. 4d**), consistent with the preserved cytotoxicity and anti-tumour response. In parallel, the frequency of CD4⁺CD25⁺Foxp3⁺ regulatory T cells (Tregs) were reduced (**Fig. 4e**) after co-administration of pLNP^CAR^ and LNP^IL-7^, indicating a less immunosuppressive TME.

Phenotypic analysis of CD8⁺ T cells further showed a drastic reduction in terminally exhausted PD-1⁺TCF-1⁻Tim-3⁺ (Tex) cells, accompanied by an increase in PD-1⁺TCF-1⁺Tim-3⁻CXCR5⁺CD127⁺ (Tpex) cells, in mice receiving co-administration of pLNP^CAR^ and LNP^IL-7^ compared with pLNP^CAR^ alone (**Fig. 4f, g**). As Tex cells reflect terminal dysfunction, whereas Tpex cells retain more stem-like, self-renewing phenotype associated with sustained antitumour immunity, this shift is consistent with the Tscm-biased state observed after IL-7 co-treatment. It may contribute to the improved durability of the antitumour response. A similar shift toward increased Tpex and reduced Tex cells was also observed in secondary lymphoid tissues, lymph nodes and spleen (**Supplementary Fig. 4a-d**). In addition, immunofluorescence analysis revealed increased CD86⁺ antigen-presenting cells within the tumour after co-administration of pLNP^CAR^ and LNP^IL-7^ (**Fig. 4h**), suggesting a shift toward a more immunostimulatory polarization state, accompanied by increased tumour apoptosis (**Supplementary Fig. 4e**).

The increased T cell stemness and reduced exhaustion led to enhanced anti-tumour effects. IVIS imaging of tumours (**Fig. 4i**) showed that in vivo pLNP^CAR^ treatment combined with LNP^IL-7^ elicited a more durable and robust antitumour response than pLNP^CAR^ alone (**Fig. 4i-l**), extending survival to 42 days, compared with 35 days in mice receiving pLNP^CAR^ alone (**Fig. 4m**).

### Combined pLNP^CAR^, LNP^IL-7^, and anti-PD-1 therapy controls established DLL3 SCLC tumours

Given that co-administration of pLNP^CAR^ and LNP^IL-7^ increased the frequency of intratumour Tpex cells, and recent studies have indicated that Tpex populations are the predominant responders to PD-1 immune checkpoint blockade (ICB)^31–34,40^, we next evaluated whether this shift could enhance responsiveness to aPD-1 therapy.

In an established subcutaneous LLC-DLL3-Luc tumour model, treatment was initiated on day 6 after tumour inoculation (**Fig. 5a**). To demonstrate the synergistic effects better, we reduced the LNP administration from 3 to 2 doses. Coadministration of pLNP^CAR^, LNP^IL-7^ and aPD-1 significantly improved antitumour effect and survival, achieving complete responses in 5 of 6 mice, whereas pLNP^CAR^ alone mediated complete responses in 2 of 6 mice (**Fig. 5b-c**). Notably, the triple-combination regimen also outperformed the three-dose LNP treatment control (data previously presented in **Fig. 2o** reused here for comparison, shown as grey dashed lines), further highlighting the therapeutic synergy of IL-7 supplementation and PD-1 blockade. No obvious treatment-related body weight loss was observed in any group (**Fig. 5d**). Histopathological examination by H&E-stained sections revealed no evident tissue damage in major organs. Notably, the spleen displayed marked immune activation, consistent with enhanced systemic immune engagement (**Supplementary Fig. 5a**). These findings together support potent and well-tolerated antitumour activity of this combinatorial immunotherapy against established tumours.

**Fig. 5.**
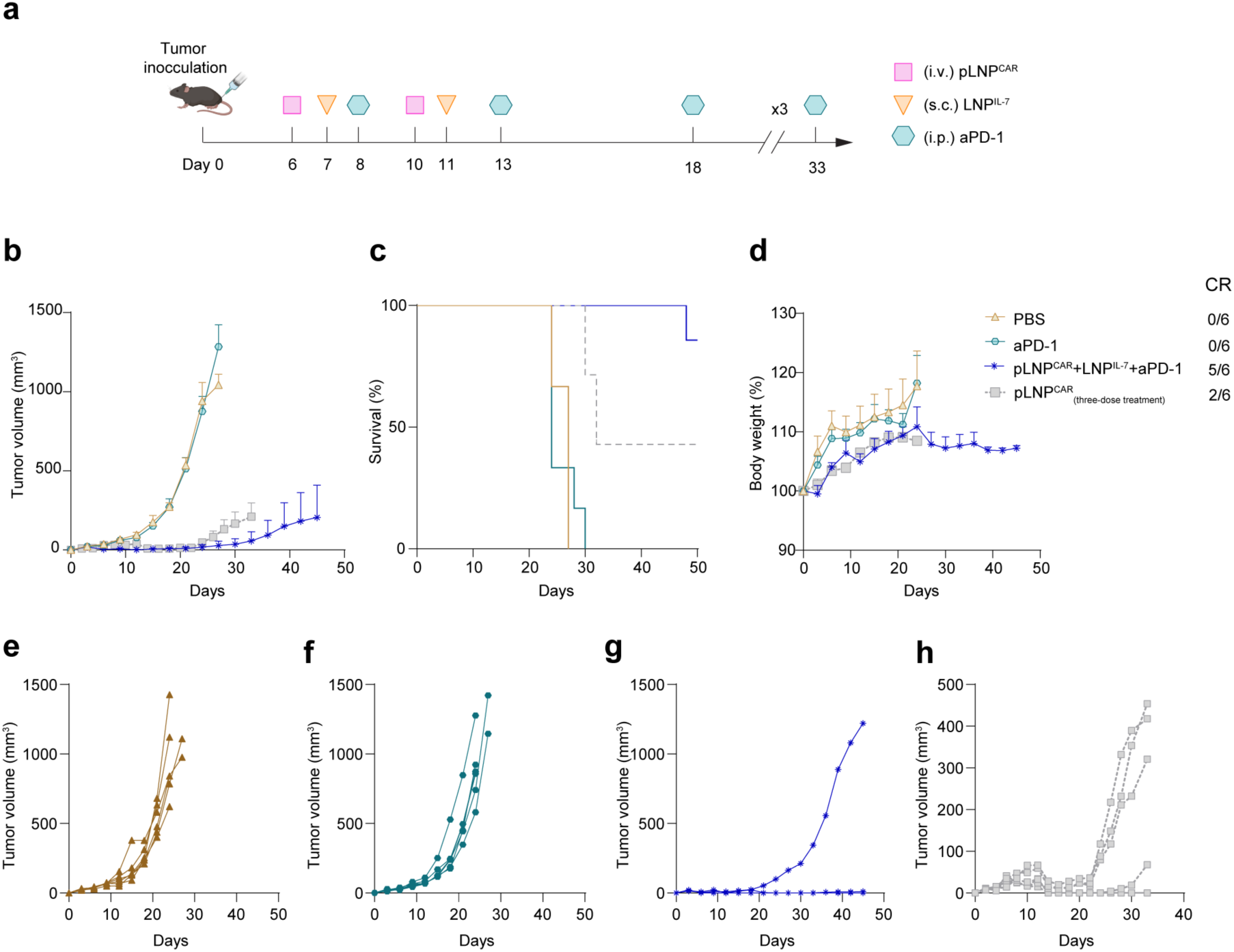
Co-administration of pLNP^CAR^ and LNP^IL-7^ synergises with anti-PD-1 checkpoint blockade to eradicate established SCLC subcutaneous tumour. **a**, Experimental timeline. Mice bearing subcutaneous DLL3⁺ SCLC tumour received intravenous pLNP^CAR^ at indicated time points, followed by subcutaneous LNP^IL-7^ administration and intraperitoneal anti–PD-1 antibody. **b**, Tumour growth kinetics. Combination therapy (pLNP^CAR^ + LNP^IL-7^ + aPD-1) markedly suppressed tumour progression compared with PBS, anti–PD-1 alone, or pLNP^CAR^ monotherapy. **c**, survival rate showing significantly prolonged survival and durable responses in the triple combination group. **d**, Body weight changes normalized to body weight on day 0 (mean ± s.e.m). **e–h**, Individual tumour growth curves for each group: PBS (**e**), aPD-1 (**f**), pLNP^CAR^ + LNP^IL-7^ + aPD-1 (**g**), and pLNP^CAR^ (data previously presented in Fig. 2o reused here for comparison showed as grey dashed lines) (**h**).

We next evaluated therapeutic efficacy in an orthotopic SCLC_DLL3 tumour model. Under a two-dose CAR-T regimen (**Fig. 6a**), co-administration of pLNP^CAR^, LNP^IL-7^, and aPD-1 significantly prolonged survival to day 49 (**Fig. 6b-d**). Analysis of tumour microenvironment revealed evident remodelling, characterized by a reduction in pan-cytokeratin tumour regions with increased infiltration of CD45^+^ leukocytes (**Fig. 6e**). H&E and CD3 immunostaining further supported reduced viable tumour burden and increased T-cell accumulation within the tumour (**Fig. 6e**). Collectively, these results demonstrate that pLNP-mediated in vivo CAR-T therapy combined with LNP^IL-7^ and aPD-1 blockade effectively remodels the tumour microenvironment and confers durable survival benefit.

**Fig. 6.**
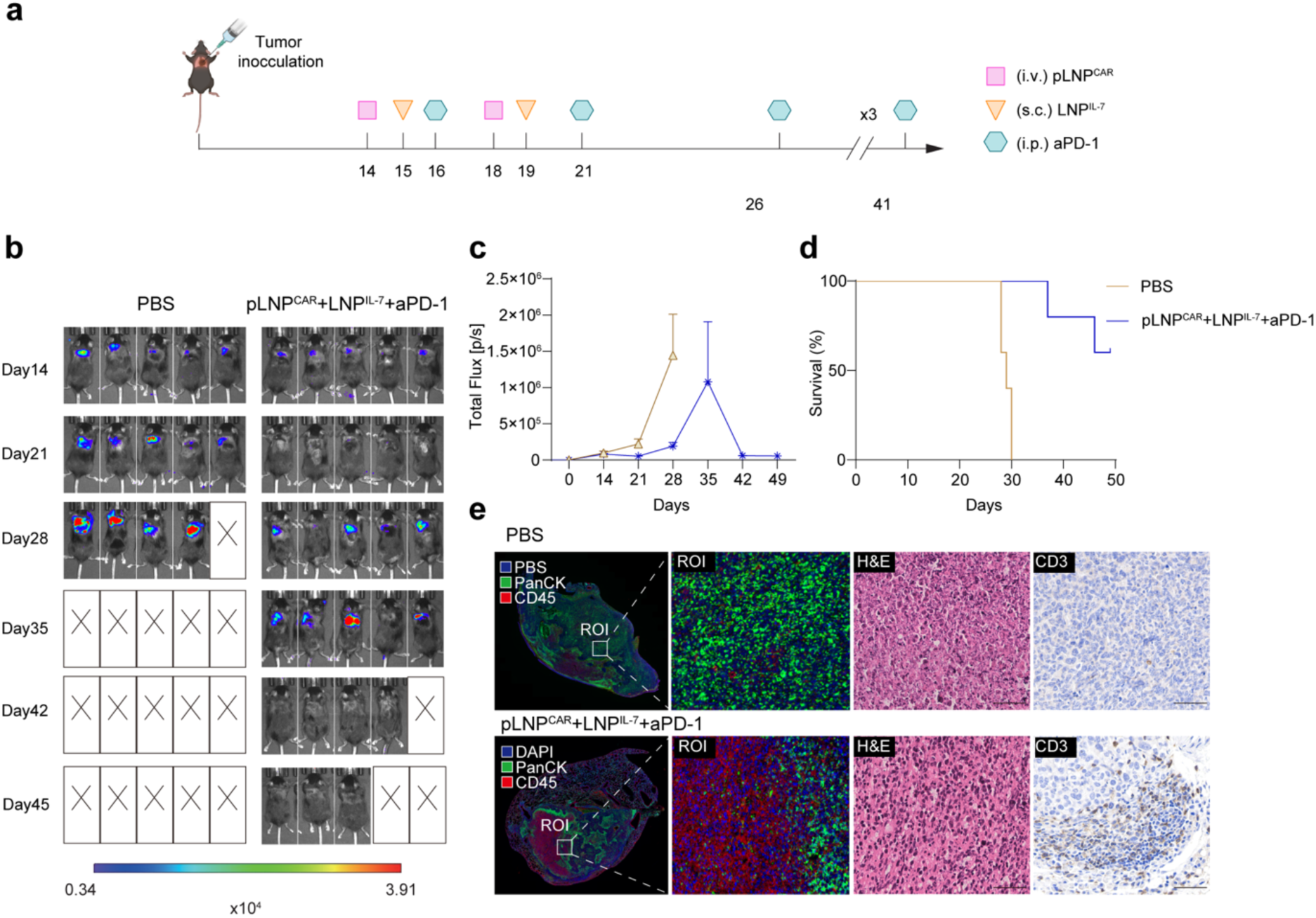
In vivo CAR T therapy combined with IL-7 and aPD-1 checkpoint blockade induced durable tumour regression and prolonged survival in a orthotopic tumour model. **a**, Experimental timeline; Mice bearing established orthotopic DLL3⁺ SCLC tumours received intravenous pLNP^CAR^ followed by subcutaneous LNP^IL-7^ and repeated intraperitoneal anti–PD-1 administration at the indicated time points. **b**, Bioluminescence image of tumour-bearing mice. **c**, tumour growth kinetics. **d**, survival rate. **e**, Regions of interest from mice tumour, immunofluorescence, H&E staining, and CD3 immunohistochemistry showing reduced tumour burden and enhanced T-cell infiltration in treated tumours, consistent with tumour microenvironment remodelling.

**Fig. 7.**
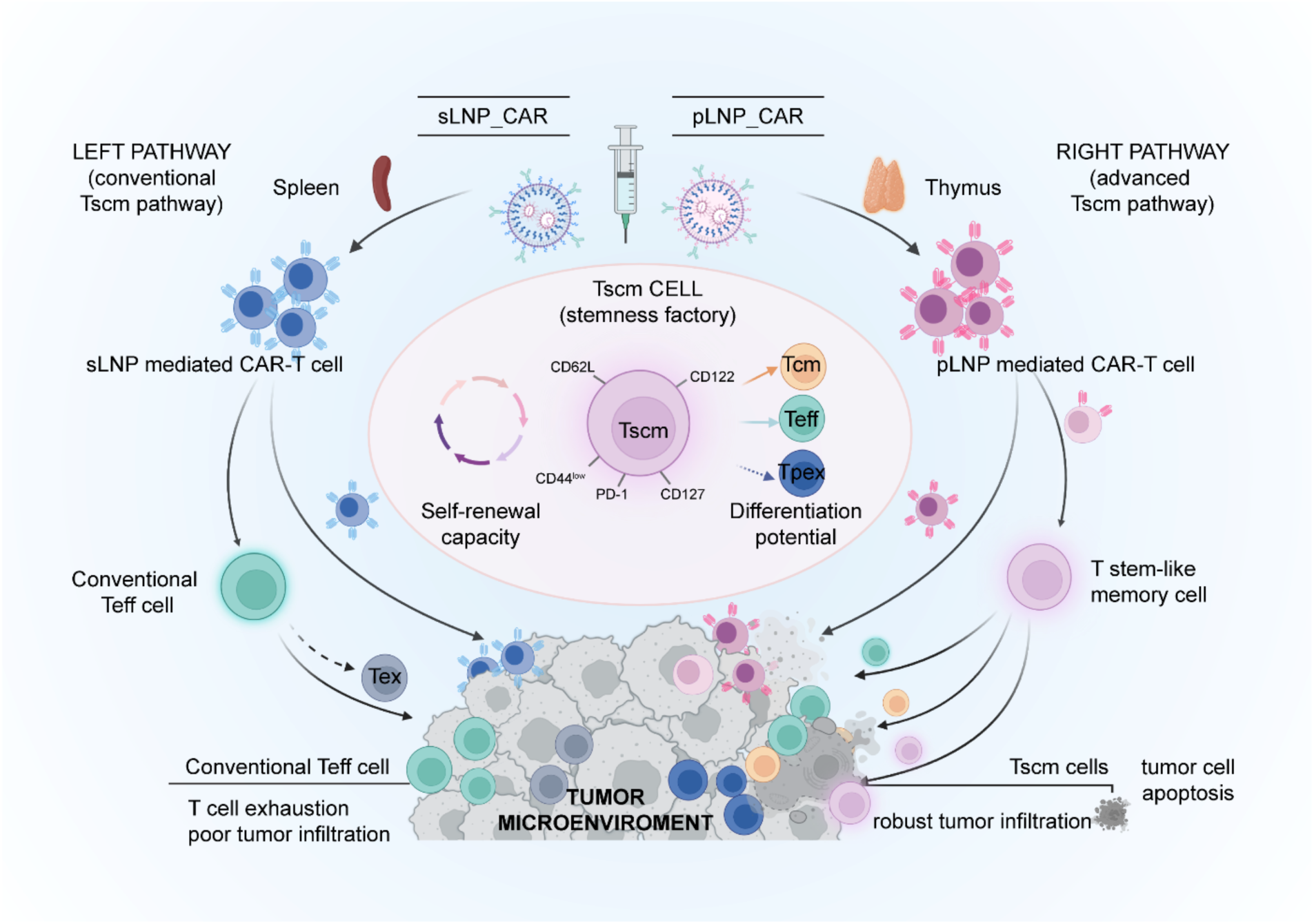
Schematic Illustration of thymus-targeted lipid nanoparticles generate stem-like memory CAR T cells in vivo for durable anti-tumour immunity. Left pathway; conventional splenic targeting (sLNP_CAR) yields effector CAR T cells (Teff) prone to rapid exhaustion (Tex) and poor tumour infiltration. Right pathway; thymus-targeted delivery (pLNP_CAR) predominantly generates stem-like memory CAR T cells (Tscm). These Tscm cells possess self-renewal capacity and multipotent differentiation potential, promoting robust tumor infiltration with tumor microenvironment remodeling for durable and potent anti-tumour responses.

## Discussion

In this study, we established a lymphoid-organ-selective LNP platform for in vivo CAR-T programming. Through systematic formulation screening, we identified pLNP as a formulation with preferential delivery to the thymus and lymphoid tissues. After antibody conjugation, pLNP enabled efficient CAR programming in T cells while preserving this organ-level tropism. Compared with sLNP, pLNP generated CAR-T cells enriched in naïve and stem-like memory phenotypes and supported prolonged CAR persistence, which translated into superior antitumour efficacy in both subcutaneous and orthotopic DLL3-positive SCLC models. These findings support lymphoid-selective in vivo CAR-T programming as a strategy to improve CAR-T durability in solid tumours.

A key finding of this study is that pLNP and sLNP differed not only in tissue tropism but also in the cellular states of the CAR-T cells they generated. pLNP preferentially engaged thymic and other lymphoid T-cell compartments, producing CAR-T cells enriched in Tn and Tscm populations, whereas sLNP showed a stronger bias toward peripheral blood and a more effector-like phenotype (**Fig. 2f-k**). In this setting, the benefit of pLNP is therefore not simply more CAR delivery, but the generation of a less differentiated CAR-T pool associated with longer persistence. This distinction is particularly important in solid tumours, where therapeutic durability depends not only on initial cytotoxicity but also on the maintenance of long-lived and renewable T-cell states. At the same time, although our data clearly establish the thymic/ lymphoid bias of pLNP, they do not yet define the delivery mechanism of this organ selectivity. Ongoing analysis of the *in vivo* protein corona may help explain how formulation-specific physicochemical features contribute to this targeting behaviour.

IL-7 is a central cytokine for T-cell survival, homeostasis, and the maintenance of less differentiated T-cell states. It has also been associated with improved persistence and functional fitness of engineered T cells. In our system, co-administration of LNP^IL7^ with pLNP^CAR^ enhanced CAR persistence in peripheral blood, increased Tscm frequencies in lymph nodes and spleen, and augmented cytotoxicity, including granzyme B and transient IL-2 production. Importantly, these effects were not accompanied by overt inflammatory amplification, as IL-6 remained low (**Fig. 3e-h**). Together, these findings suggest that IL-7 acts in this setting primarily by reinforcing CAR-T persistence and functional durability, rather than by broadly amplifying inflammatory activation.

The immunosuppressive tumour microenvironment remains a major barrier to durable CAR-T efficacy in solid tumours. In this setting, co-administration of pLNP^CAR^ and LNP^IL7^ reinforced CAR-T persistence and stem-like features in lymphoid tissues and reshaped the intratumoural immune landscape. The frequency of immunosuppressive T regulatory cells has decreased. Remarkably, CD8⁺ T-cell states shifted away from terminally exhausted Tex cells toward progenitor exhausted Tpex cells, accompanied by increased granzyme B expression (**Fig. 4k-l**). Recent studies have highlighted Tpex cells as a stem-like exhausted subset that retains self-renewal capacity and sustains antitumour immunity under chronic antigen exposure, making this population particularly relevant to durable CAR-T responses in solid tumours^30–31,40,52^. As a result, the increase in Tpex cells after LNP^IL7^ co-administration is consistent with a more durable and therapeutically responsive CAR-T state. These findings further suggest that reinforcement of stem-like CAR-T features outside the tumour can translate into improved functional quality within the tumour. Additional evidence of a less suppressive and more immunostimulatory microenvironment was provided by increased CD86-positive antigen-presenting cells and increased tumour apoptosis in combined therapy (**Fig. 4m**). These findings further suggest that reinforcement of stem-like CAR-T features in secondary lymphoid tissues may translate into improved intratumoural CAR-T fitness and more favourable tumour microenvironmental remodelling.

The increase in Tpex cells after LNP^IL-7^ reinforcement also provided a strong rationale for combining this platform with immune-checkpoint blockade treatment, since recent studies have shown that Tpex cells are the major responders to PD-1 immune checkpoint blockade^31–40^. Therefore, triple therapy with pLNP^CAR^, LNP^IL7^, and aPD-1 produced potent and durable antitumour responses, yielding superior tumour control and prolonged survival benefit in both established subcutaneous and orthotopic DLL3 SCLC tumour models (**Fig. 5**, **Fig. 6**). Together, these findings demonstrate that the combined thymus-oriented pLNP^CAR,^ LNP^IL7^, and aPD-1 blockade strategy enables more persistent in vivo CAR-T programming by sustaining a more stem-like CAR-T state, thereby conferring robust and prolonged antitumour activity. More broadly, this study underscores the potential of LNP-based platforms to precisely engineer immune responses in situ by integrating organ-selective delivery with rational immunomodulation.

Several limitations of this study should be considered. Although pLNP showed clear thymic and lymphoid selectivity, the molecular determinants underlying this organ tropism remain incompletely defined. In addition, while pLNP delivery was associated with Tn/Tscm enrichment and prolonged CAR persistence, the present data do not directly establish how thymic targeting influences the developmental trajectory of the resulting CAR-T cells. Similarly, although LNP^IL-7^ co-administration shifted the balance from Tex toward Tpex and improved responsiveness to PD-1 blockade, the mechanistic links among cytokine reinforcement, T-cell state transitions, and checkpoint sensitivity require further clarification. Future studies should therefore define the uptake mechanisms governing pLNP organ selectivity, more directly trace the developmental origin of pLNP-generated CAR-T cells and dissect how cytokine reinforcement and checkpoint blockade interact over time within distinct tumour contexts.

## Methods

### Materials

ALC0315 was purchased from AVT (Shanghai, China). LP01 and 1,2-distearoyl-sn-glycero-3-phosphoethanolamine-N-[maleimide(polyethylene glycol)-2000] (ammonium salt) (DSPE-PEG (2000) Maleimide) was purchased from Avanti Research (U.S.). GT7 and OT13 refer to patents (WO2025256311A1) and (WO2024094098A1), respectively, which were synthesized via WUXI APP TEC (China). 1, 2 - Dioctadecanoyl- sn-glycero- 3 - phophocholine (DSPC), dioleoylphosphatidylethanolamine (DOPE), cholesterol, 1,2-distearoyl-sn-glycero-3-phosphoethanolamine-N-[amino(polyethylene glycol)-2000] and 1,2-dimyristoyl-rac-glycero-3-methoxypolyethylene glycol-2000 (DMG-PEG2000) were purchased from AVT (Shanghai, China). N-succinimidyl S-acetylthioacetate (SATA) was purchased from Thermo Fisher (U.S.) Dimthyl sulfoxide (DMSO) and hydroxylamine were purchased from MCE (Shanghai, China). Purified anti-mouse CD5 Antibody was purchased from BioLegend (China).

### Mice

C57BL/6J were purchased from Beijing Sinogenetic Biotechnology. Ai9 mice were purchased from Kexing. All animal experiments complied with the standards of the Institutional Animal Care and Use Committee of the Institute of Process Engineering, Chinese Academy of Sciences (approval ID: IPEAECA2024039).

### RNA synthesis and Lipid nanoparticle formulation

Luc, Enhanced-GFP, and Cre recombinase mRNA constructs encoding fragments of each sequence were cloned into pUC57-Kan vectors to generate plasmids (Nanjing GenScript Biotechnology Co., Ltd.). Briefly, a T7 promoter, the gene of interest (GOI), optimized 5’(3’)-untranslated regions and poly-A tails. Plasmids were extracted using Quantum Prep Plasmid Midiprep kit (BioRad, 7326120). mRNA was in vitro transcribed and capped using a T7 Co-transcription RNA synthesis kit (SuZhou Novoprotein Scienticic Co.Ltd.). DLL3 encoded circular RNA was provided by Thoracic Oncology Institute, Peking University People’s Hospital (Beijing, China).

A library of 48 LNPs was formulated using an N:P ratio of 4.5 as previously described^19^ lipid to nucleic acid with an ionizable lipid and excipient formulation (**Supplementary Fig. 1a**). Briefly, LNPs were prepared via microfluidic mixing of an aqueous solution of the mRNA and an ethanolic solution of the lipid components at a ratio of 3:1 for all LNPs. The ethanol phase consists of cationic lipid, 1,2-distearoyl-sn-glycero-3phosphocholine (DSPC) or dioleoylphosphatidylethanolamine (DOPE), cholesterol, and polyethylene glycol (DMG-PEG2000) or 1,2-distearoyl-sn-glycero-3-phosphoethanolamine-N-[amino(polyethylene glycol)-2000 (DSPE-PEG2000). For antibody-conjugated LNP, 0.5% ratio of PEG was replaced with maleimide (polyethylene glycol-2000) (DSPE-PEG(2000) maleimide). A separate aqueous phase containing mRNA was prepared in 10mM citrate buffer (pH=4).

Antibody-conjugated LNPs were performed using SATA–maleimide chemistry^12–16^. Purified rat anti-mouse CD5 antibody was first thiolated by reaction with SATA (N-succinimidyl S-acetylthioacetate) according to the manufacturer’s protocol, followed by deacetylation with hydroxylamine to expose free sulfhydryl groups. Separately, freshly prepared LNP–mRNA formulations containing maleimide-functionalized PEG–lipids were equilibrated at room temperature. The thiolated antibody was then added to the maleimide-bearing LNPs at a defined molar ratio and incubated under gentle agitation for 1–2 h to allow covalent thioether bond formation. Unreacted antibody was removed by ultrafiltration, and the resulting antibody-conjugated LNP–mRNA particles were buffer-exchanged into PBS and stored at 4 °C until use.

The particle size of LNP was characterized via dynamic light scattering (DLS). Briefly, LNPs were diluted 200X in 1xPBS solution. Z-average diameter and polydispersity index were recorded using a Zetasizer Nano (Malvern). RNA encapsulation efficiencies of each LNPs were measured using a Quant-iT RiboGreen RNA Quantification kit (Thermo Fisher). The fluorescence intensity was measured under the condition of excitation light with a wavelength of 480nm and emission light with 520nm using a microplate reader. The fluorescence intensity of the sample was converted into the corresponding mRNA concentration in the standard curve. Encapsulation efficiency formula = (total mRNA concentration - free mRNA concentration) / total mRNA concentration of demulsified sample ×100%.

### In vivo biodistribution studies

#### In vivo luciferase mRNA LNP delivery

LNPs encapsulating luciferase mRNA were administered i.v. to wild-type 8-week-old male C57BL6/J mice at doses of 0.5 mg kg^−1^ of mRNA. Six hours following the administration, luminescence imaging was performed using Living Image software on an IVIS (PerkinElmer). Five minutes before imaging, 200 µL (15mg/mL) D-luciferin potassium salt (APOxBIO) was i.p. administered to mice. Mice were then euthanized with CO2, and the heart, lungs, liver, kidneys, spleen, lymph node, and thymus were dissected and imaged. To quantify luminescence, Living Image Software (PerkinElmer) was used to place a rectangular region of interest. The spleen-to-liver ratio was calculated by dividing the luminescence in the spleen by the luminescence in the liver.

### In vivo Cre mRNA-LNP delivery

LNPs encapsulating Cre recombinase mRNA were administered i.v. to AI9 reporter mice at a dose of 1.5 mg kg^−1^ of mRNA^19^. Five days after the administration, peripheral blood, liver, spleen, lymph node, and thymus were dissected. For flow cytometry, spleen, lymph node, and thymus were mechanically ground to a single-cell suspension. For immunofluorescence sectioning, Liver, spleen, lymph node, and thymus were fixed with 4% paraformaldehyde overnight and then dehydrated with a 30% glucose solution.

### In vivo DLL3 CAR T cell generation

LNPs encapsulating DLL3 encoded mRNA were administered i.v. to wild-type C57 at doses of 0.5, 1, 1.5 mg kg-1of mRNA. Mice were sacrificed after 6H, 24H, 48H, 72H, 96H, 168H, 14D, 21D, and 28D administration of LNP. Peripheral blood, spleen, lymph node, and thymus were made into single-cell suspensions for flow cytometry.

### Flow cytometry

Beckman Coulter flow cytometry (CytoFLEX LX) was used in this study. Tumour or immune organs (mentioned above) were mechanically ground and passed through a 70 µm strainer to create single-cell suspensions. Red blood cells were removed by incubation with RCL according to the manufacturer’s instructions. Then, cells were washed and resuspended in MACS buffer (containing 0.5% BSA, 10mM EDTA in 1x PBS). Cells were then incubated with antibodies for 30 minutes at 4 °c in the dark. The following antibody were used: PerCP/ Cyanine5.5 anti-mouse CD45 antibody (BioLegend, clone 30-F11, 103132), APC anti-mouse CD3 antibody (BioLegend, clone 17A2, 100236), APC/Cyanine7 anti-mouse CD4 antibody (BioLegend, clone RM4-5, 100526), Pacific Blue anti-mouse CD8a antibody (BioLegend, clone 53-6.7), Alexa Fluor 700 anti-mouse CD19 antibody (BioLegend, clone 6D5, 115528), APC anti-mouse F4/80 antibody (BioLegend, clone BM8, 123116), Alexa Fluor 488 anti-mouse/human Cd11a antibody (BioLegend, clone M1/70, 101217), Pacific Blue anti-mouse CD11c antibody (BioLegend, clone N418, 117322), PE/Cyanine 7 anti-mouse/human CD44 antibody (BioLegend, clone IM7, 103030), Brilliant Violet 711 anti-mouse CD62L antibody (BioLegend, clone MEL-14, 104445), FITC anti-mouse CD69 antibody (BioLegend, clone H1.2F3), Brilliant Violet 605 anti-mouse PD-1 (BioLegend, clone 29F.1A12, 135220), PE anti-mouse CD69 antibody (BioLegend, clone H1.2F3, 104507), TCF1/TCF7 (C63D9) Rabbit mAb (APC conjugate) (BD Phaemingen, 37636S), PE anti-mouse CD127 (IL-7rα) antibody (BioLegend, clone A7R34, 135009), PE/Cyanine 7 anti-mouse CXCR5 (BioLegend, clone L138D7, 145516), PerCP/Cyanine 5.5 anti-mouse Tim-3 (BioLegend, clone RMT3-23, 119718), PerCP/Cyanine5.5 anti-mouse CD122 (IL-2rβ) antibody (BioLegend, clone TM-B1, 123211), FITC anti-mouse Ly-6A/E (Sca-1) antibody (BioLegend, clone D7, 108105), PE anti-mouse IL-2 antibody (BioLegend, clone JES6-5H4, 503807), APC anti-mouse CD25 antibody (BioLegend, clone PC61, 1012011), PE-Cyanine 7 Foxp3 anti-mouse antibody (Thermo Fisher, clone FJK-16s, 25-5773-82), APC anti-mouse IFN-γ antibody (BioLegend, clone XMG1.2, 505810), PE anti-mouse TNF-α antibody (BioLegend, clone MP6-XT22, 506306), PE/Cyanine 7 anti-human/ mouse Granzyme B antibody (BioLegend, clone QA18A28, 396410). PE MonoRab Rabbit Anti-scFv Cocktail (Nanjing GenScript Biotechnology Co., Ltd., A02285) were used to analysis CAR expression and Ghost Dye Violet 510 (Cytek, 13-0870) were used for determining cell viability in this paper.

### In vivo studies of mice tumour model

To evaluate in vivo anti-tumour efficiency, mice were housed with a 12 h light and 12 h dark cycle at a temperature range of 20-24 °C and humidity between 30-70%. For subcutaneous tumour studies, 1.0 x 10^^6^ LLC_DLL3_Luc tumour cells were injected on the left side of the back of each male C57BL/6J mouse on day 0. Five days after tumour inoculation, LNP were intravenously injected through the tail vein at a dose of 1.5 mg kg-1 on day 6, day 10 or day 12 for sLNP and pLNP. For combination therapy, LNP encapsulating IL7_mRNA was subcutaneously injected (10 ug each) 24h after pLNP. Anti-mouse PD-1 (Bio Xcell) were intraperitoneally injected (100 ug each) starting at 2 days after pLNP administration with 5 days interval. Tumour size was measured with callipers every 3 days (2 days for Fig. 2), calculated as ellipsoids (V=length x width^2 x ½). Mice were euthanized when tumour volume had exceeded 1000mm^3^ or tumour ulceration of more than 5mm in diameter had been observed.

For the orthotopic solid tumour study, mice were completely anaesthetized. On the left side of the back of each male C56BL/6J mouse, a small incision was made to access the thoracic cavity, followed by injection of 2.0 × 10^^6^ LLC_DLL3_LUC tumour cells suspended in 100ul of PBS into the lung parenchyma. The incision was closed with an appropriate suture to prevent leakage and contamination. Tumours were allowed to grow for approximately 14 days. When tumour burden reaches 1×10^^5^ flux via bioluminescence imaging (PerkinElmer). The experiment group received tail vein treatment as illustrated in the experimental timeline and figure legend. Mice were euthanized when bioluminescence exceeded the maximum values or mice exhibited clear signs of distress, such as pronounced loss of appetite, lethargy or failure to thrive. In the histology study, tumours were collected 3 days after the treatments.

### Histology

Subcutaneous and orthotopic tumour-bearing C57BL/6J mice tumours and organs were collected 3 days after two doses of pLNP^CAR^ or pLNP^CAR^ and LNP^IL-7^. Tissues were fixed with 4% paraformaldehyde solution in PBS overnight at 4 °C. Formalin-fixed, paraffin-embedded tissue samples were sectioned, followed by H&E and immunohistochemistry staining. Antigen and slide imaging were provided by Solrbio Science & Technology Co.,Ltd. (Beijing, China).

### Statistical analysis

Experimental data were analysed using GraphPad Prism version 10.0 and ImageJ software. Group differences were evaluated using unpaired, two-tailed Student’s t-tests or two-way ANOVA. Statistical significance levels were defined as p-values indicated in the figures. Data are presented as means ± SEM. All experiments were performed with at least three replicates per group (n ≥ 3).

## Acknowledgements

We thank Dr Mantang Qiu, Dr Jingsheng Cai (Peking University People’s Hospital, Beijing, China) for generously providing the RNA materials used in this study. This work was supported by the Strategic Priority Research Program of the Chinese Academy of Sciences (XDB1310000). National Key Research and Development Program of China (2021YFC2302400 to Y.X., 2021YFC2302605 to G. M., 2024YFC2310700 to X. H.), the Major Program of the National Natural Science Foundation of China (T2394503 to Y.X., T2394501 to G. M.). Additional support was provided by the CAS Project for Young Scientists in Basic Research (YSBR-010 to Y.X.), and the IPE Project for Frontier Basic Research (QYJC-2023-04 to Y.X.).

## References

1. Maude, S. L. et al. Chimeric antigen receptor T cells for sustained remissions in leukemia. N. Engl. J. Med. 371, 1507–1517 (2014).

2. Schuster, S. J. et al. Chimeric antigen receptor T cells in refractory B-cell lymphomas. N. Engl. J. Med. 377, 2545–2554 (2017).

3. June, C. H., O’Connor, R. S., Kawalekar, O. U., Ghassemi, S. & Milone, M. C. CAR T cell immunotherapy for human cancer. Science 359, 1361–1365 (2018).

4. Gattinoni, L. et al. Acquisition of full effector function in vitro paradoxically impairs the in vivo antitumour efficacy of adoptively transferred CD8⁺ T cells. J. Clin. Invest. 115, 1616–1626 (2005).

5. Fraietta, J. A. et al. Determinants of response and resistance to CD19 CAR T cell therapy. Nat. Med. 24, 563–571 (2018).

6. Langer, R. & Weissman, D. mRNA vaccines — a new era in vaccinology. Nat. Rev. Drug Discov. 17, 261–279 (2018).

7. Sahin, U., Karikó, K. & Türeci, Ö. mRNA-based therapeutics — developing a new class of drugs. Nat. Rev. Drug Discov. 13, 759–780 (2014).

8. Rurik, J. G. et al. CAR T cells produced in vivo to treat cardiac injury. Science 375, 91–96 (2022).

9. Parayath, N. N. et al. In situ programming of CAR T cells using synthetic DNA nanocarriers. Nat. Nanotechnol. 15, 996–1003 (2020).

10. Billingsley, M. M. et al. Ionizable lipid nanoparticle-mediated mRNA delivery for human CAR T cell engineering. Nat. Biomed. Eng. 4, 933–942 (2020).

11. Sterner, R. C. & Sterner, R. M. CAR-T cell therapy: current limitations and potential strategies. Blood Cancer J. 11, 69 (2021).

12. Schmid, D. et al. T cell-targeting nanoparticles focus delivery of immunotherapy to improve antitumour immunity. Nat. Commun. 8, 1747 (2017).

13. Howard, M. D., Jay, M., Dziubla, T. D. & Lu, X. PEGylated liposomes conjugated with antibodies using SATA–maleimide chemistry. Mol. Pharm. 11, 2262–2270 (2014).

14. Marcos-Contreras, O. A. et al. Vascular targeting of nanocarriers using antibody–liposome conjugates prepared via SATA–maleimide chemistry. J. Control. Release 201, 37–45 (2015).

15. Tombácz, I. et al. Targeted lipid nanoparticles for mRNA delivery via antibody conjugation using thiol–maleimide chemistry. Mol. Ther. 29, 3293–3304 (2021).

16. Geisler, Hannah C., et al. “Preparation of targeted lipid nanoparticles for precision nucleic acid delivery.” Nature Protocols 1–36 (2026).

17. Álvarez-Benedicto, E. et al. Spleen SORT LNP generated in situ CAR T cells extend survival in a mouse model of lymphoreplete B cell lymphoma. Angew. Chem. Int. Ed. 62, e202310395 (2023).

18. Cheng, Q., Wei, T., Farbiak, L. et al. Selective organ targeting (SORT) nanoparticles for tissue-specific mRNA delivery and CRISPR–Cas gene editing. Nat. Nanotechnol. 15, 313–320 (2020).

19. Gao, W. et al. Resolving the mRNA encapsulation–release trade-off via compensatory forces in engineered ionizable lipids. Adv. Mater. 37, 2409483 (2025).

20. Cai, Jingsheng, et al. Engineered circular RNA-based DLL3-targeted CAR-T therapy for small cell lung cancer. Experimental hematology & oncology 14.1 35 (2025).

21. Zhang, R. et al. Helper lipid structure influences protein adsorption and delivery of lipid nanoparticles to spleen and liver. Biomater. Sci. 9, 1673–1684 (2021).

22. LoPresti, S. T. et al. The replacement of helper lipids with charged alternatives in lipid nanoparticles facilitates targeted mRNA delivery to the spleen and lungs. J. Control. Release 345, 319–331 (2022).

23. Kranz, L. M. et al. Systemic RNA delivery to dendritic cells exploits antiviral defence for cancer immunotherapy. Nature 534, 396–401 (2016).

24. Newick, K., O’Brien, S., Moon, E. & Albelda, S. M. CAR T cell therapy for solid tumours. Annu. Rev. Med. 68, 139–152 (2017).

25. Binnewies, M. et al. Understanding the tumour immune microenvironment. Nat. Med. 24, 541–550 (2018).

26. Joyce, J. A. & Fearon, D. T. T cell exclusion, immune privilege, and the tumour microenvironment. Science 348, 74–80 (2015).

27. Wherry, E. J. T cell exhaustion. Nat. Immunol. 12, 492–499 (2011).

28. Gattinoni, L., Speiser, D. E., Lichterfeld, M. & Bonini, C. T cell persistence in adoptive immunotherapy. Nat. Rev. Cancer 17, 451–467 (2017).

29. Schietinger, A. & Greenberg, P. D. Tolerance and exhaustion: defining mechanisms of T cell dysfunction. Trends Immunol. 35, 51–60 (2014).

30. Liu, Qiao, et al. “Tumor-specific memory CD8+ T cells are strictly resident in draining lymph nodes during tumorigenesis.” Cellular & molecular immunology 20.4, 423–426 (2023).

31. Huang, Qizhao, et al. “The primordial differentiation of tumor-specific memory CD8+ T cells as bona fide responders to PD-1/PD-L1 blockade in draining lymph nodes.” Cell 185.22, 4049–4066 (2022).

32. Yang, Yang, et al. “Oncolytic virotherapy potentiates chemo-PD-1 immunotherapy by engaging chemo-resistant bystander CD8+ T cells.” Journal for Immunotherapy of Cancer 14.3 (2026).

33. Liu, Qiao, et al. “Chromatin landscape alteration uncovers multiple transcriptional circuits during memory CD8+ T-cell differentiation.” Protein & Cell 16.7, 575–601 (2025).

34. Im, S. J. et al. Defining CD8⁺ T cells that provide the proliferative burst after PD-1 therapy. Nature 537, 417–421 (2016).

35. Utzschneider, D. T. et al. T cell factor 1–expressing memory-like CD8⁺ T cells sustain the immune response to chronic viral infections. Immunity 45, 415–427 (2016).

36. Miller, B. C. et al. Subsets of exhausted CD8⁺ T cells differentially mediate tumour control and respond to checkpoint blockade. Nat. Immunol. 20, 326–336 (2019).

37. Pauken, K. E. et al. Epigenetic stability of exhausted T cells limits durability of reinvigoration by PD-1 blockade. Science 354, 1160–1165 (2016).

38. Beltra, J.-C. et al. Developmental relationships of four exhausted CD8⁺ T cell subsets reveal underlying transcriptional and epigenetic landscape control mechanisms. Cell 181, 1312–1326 (2020).

39. Russell, J. H. & Ley, T. J. Lymphocyte-mediated cytotoxicity. Annu. Rev. Immunol. 20, 323–370 (2002).

40. Siddiqui, I. et al. Intratumoural Tcf1⁺PD-1⁺CD8⁺ T cells with stem-like properties promote tumour control in response to vaccination and checkpoint blockade immunotherapy. Immunity 50, 195–211 (2019).

41. Klein, L., Kyewski, B., Allen, P. M. & Hogquist, K. A. Positive and negative selection of the T cell repertoire. Nat. Rev. Immunol. 14, 377–391 (2014).

42. Germain, R. N. T cell development and the CD4–CD8 lineage decision. Nat. Rev. Immunol. 2, 309–322 (2002).

43. Palmer, E. & Naeher, D. Affinity threshold for thymic selection through a T cell receptor–co-receptor zipper. Nat. Rev. Immunol. 9, 207–213 (2009).

44. Voss, J. E. et al. Thymic programming of T cell fate. Nat. Immunol. 22, 1095–1105 (2021).

45. Mazzucchelli, R. & Durum, S. K. Interleukin-7 receptor expression: intelligent design. Nat. Rev. Immunol. 7, 144–154 (2007).

46. Surh, C. D. & Sprent, J. Homeostasis of naïve and memory T cells. Nat. Immunol. 9, 13–19 (2008).

47. Mackall, C. L., Fry, T. J. & Gress, R. E. Harnessing the biology of IL-7 for therapeutic application. Nat. Rev. Immunol. 11, 330–342 (2011).

48. Pellegrini, M. et al. IL-7 engages multiple mechanisms to overcome chronic viral infection and limit organ pathology. Nat. Immunol. 12, 512–520 (2009).

49. Sportès, C. et al. Administration of rhIL-7 in humans increases in vivo TCR repertoire diversity. Blood 111, 5237–5244 (2008).

50. Beq, S. et al. IL-7 induces immunological improvement in SIV-infected rhesus macaques under antiretroviral therapy. J. Clin. Invest. 119, 997–1007 (2009).

51. Philip, M. & Schietinger, A. CD8⁺ T cell differentiation and dysfunction in cancer. Nat. Rev. Immunol. 22, 209–223 (2022).

52. Kallies, A., Zehn, D. & Utzschneider, D. T. Precursor exhausted T cells: key to successful immunotherapy? Nat. Rev. Immunol. 20, 128–136 (2020).

53. Wherry, E. J. & Kurachi, M. Molecular and cellular insights into T cell exhaustion. Nat. Rev. Immunol. 15, 486–499 (2015).

