## Supplemental Figure 1-6 for "In vivo programming of stem-like CAR T cells by lymphatic-selective lipid nanoparticle enables durable anti-tumour efficacy in orthotopic model"

**a**

| Ionizable lipid | Cholesterol | Helper lipid | Pegylated Lipid | Formulation (molar ratio) |
|---|---|---|---|---|
| ALC0315 (US10221127B2) | Cholesterol | DSPC Distearoylphosphatidylcholine | DMG-PEG2000, 1,2-dimyristoyl-rac-glycero-3-methoxypolyethylene glycol-2000 | 35:46.5:16:2.5 |
| LP-01 (US11420933B2) | | | | 40:48.5:10:1.5 |
| OT13 (WO2024094098A1) | | DOPE Dioleoylphosphatidylethanolamine | DSPE-PEG2000, 1,2-distearoyl-sn-glycero-3-phosphoethanolamine-N-[amino(polyethylene | |
| GT7 (WO2025256311A1) | | | | 45:44:9:2 |

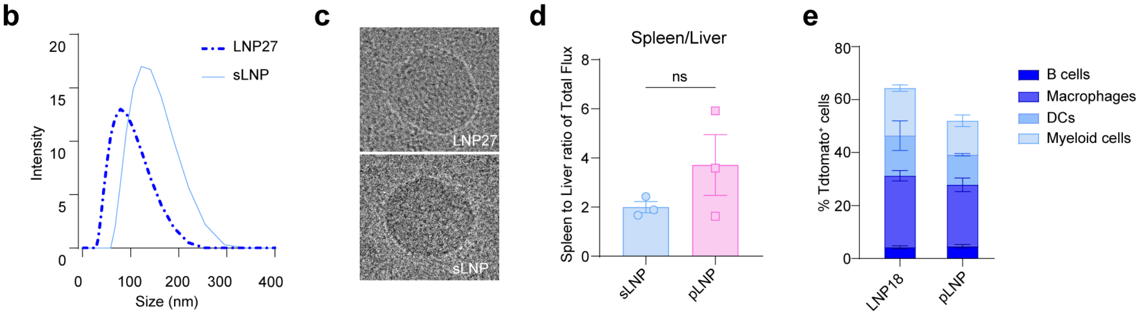


**Supplementary Fig. 1 Formulation and characterization of LNP for lymphatic targeting. a**, components of the LNP library used for in vivo screening. Four ionizable lipids (ALC0315, LP-01, OT13 and GT7) were combined with cholesterol, helper phospholipids (DSPC or DOPE) and PEG-lipids (DMG-PEG2000 or DSPE-PEG2000) at defined molar ratios to generate distinct LNP formulations. **b**, Hydrodynamic diameter distributions of representative LNP 3 and sLNP (antibody conjugated of LNP27) measured by dynamic light scattering (DLS). **c**, Representative cryogenic transmission electron microscopy (cryo-TEM) images of LNP27 and sLNP nanoparticles showing spherical morphology and uniform structural organization. **d**, Quantification of spleen-to-liver bioluminescence signal ratios after i.v. administration of sLNP and pLNP. **e,** Flow cytometric quantification of tdTomato$^{+}$ cells in lymphoid tissues following i.v. injection of LNP-encapsulated Cre mRNA in Ai9 reporter mice. Distribution of transgene expression across major immune cell populations, including B cells, macrophages, dendritic cells (DCs) and myeloid cells. Data are presented as mean ± s.e.m. Statistical significance was determined by one-way ANOVA with multiple comparisons. p values are indicated in the panels.

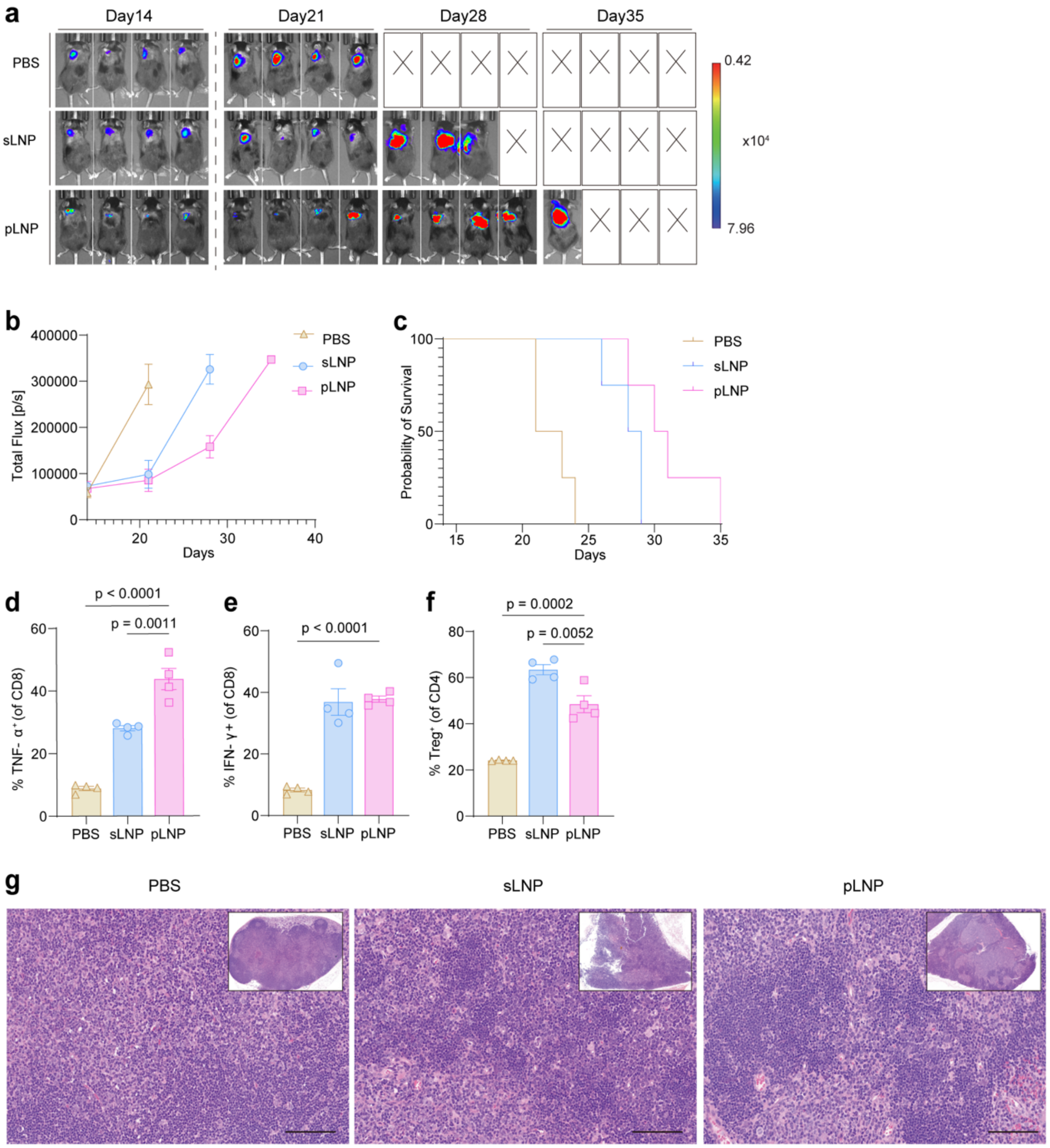


**Supplementary Fig. 2 Thymus-targeted pLNP delivery enhances antitumour immune responses in vivo. a**, Bioluminescence imaging of SCLC orthotopic tumour-bearing mice treated with PBS, sLNP, or pLNP at the indicated time points. Quantification of tumour flux (**b**), survival rate (**c**). **d-f**, Flow cytometric analysis of mice tumour. TNF-α of $CD8^+$ T cells (**d**), IFN-γ of $CD8^+$ T cells (**e**), and Treg of $CD4^+$ T cells (**f**). **g**, H&E staining of thymus showing influence of thymus differentiation following i.v. injection of sLNP or pLNP. scale bar 100μm. Data are presented as mean ± s.e.m. Statistical significance was determined by one-way ANOVA with multiple comparisons. p values are indicated in the panels.

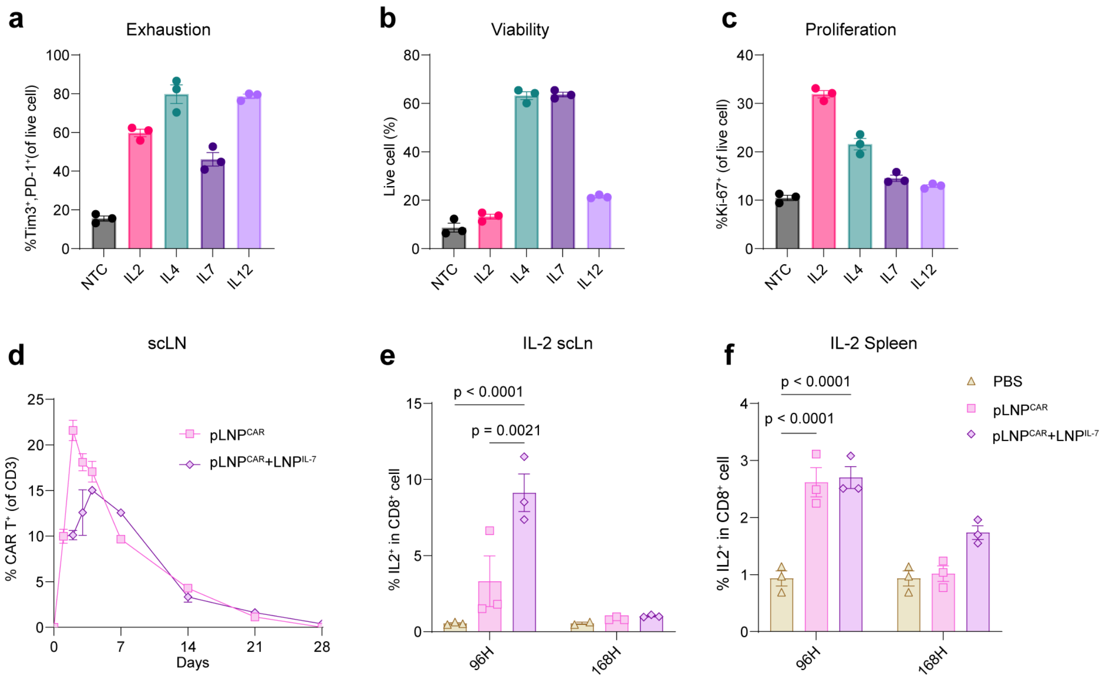


**Supplementary Fig. 3 LNP$^{IL-7}$ supports CAR T cell persistence in the lymph node and spleen. a-c**, Flow cytometry analysis of different cytokines in ex vivo T cells. Frequency of exhausted T cells (**a**), cell viability (**b**), and proliferations of T cells (**c**). **d**, Kinetics of CAR$^{+}$ T cells in lymph nodes following in vivo delivery of pLNP$^{CAR}$ or pLNP$^{CAR}$+LNP$^{IL-7}$. **e-f**, secretion of IL-2 of CD8$^{+}$ T cells at 96 h and 168 h following treatment with PBS, pLNP$^{CAR}$ or pLNP$^{CAR}$+LNP$^{IL-7}$. scLN (**e**), spleen (**f**). Data are presented as mean ± s.e.m. Statistical significance was determined by one-way ANOVA with multiple comparisons. P values are indicated in the panels.

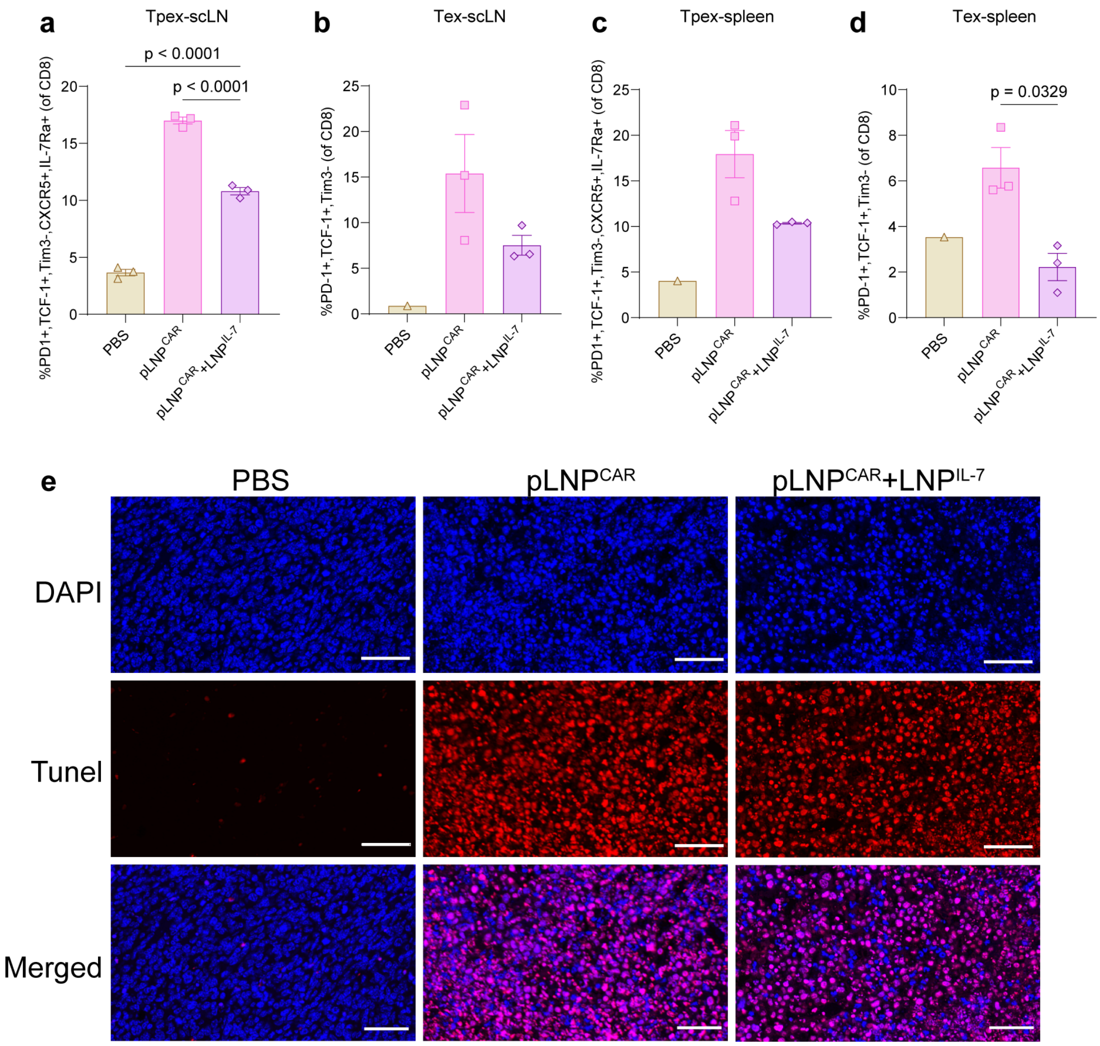


**Supplementary Fig. 4 LNP$^{IL-7}$ co-administration promotes Tpex maintenance and tumour cell apoptosis in vivo. a-d**, Flow cytometry analysis of SCLC orthotopic tumour-bearing mice spleen and lymph node. Frequency of progenitor exhausted CD8$^+$ T cells (Tpex; PD-1$^+$ TCF-1$^+$ Tim-3$^-$ CXCR5$^+$ IL-7Rα$^+$) in lymph nodes (**a**), spleen (**b**), Frequency of terminally exhausted CD8$^+$ T cells (Tex; PD-1$^+$ TCF-1$^-$ Tim-3$^+$) in lymph nodes (**c**), spleen (**d**). **e**, Immunofluorescence staining of tumour sections labelled with DAPI (blue) and TUNEL (red). Scale bars, 50 μm. Data are presented as mean ± s.e.m. Statistical significance was determined by one-way ANOVA with multiple comparisons. p values are indicated in the panels.

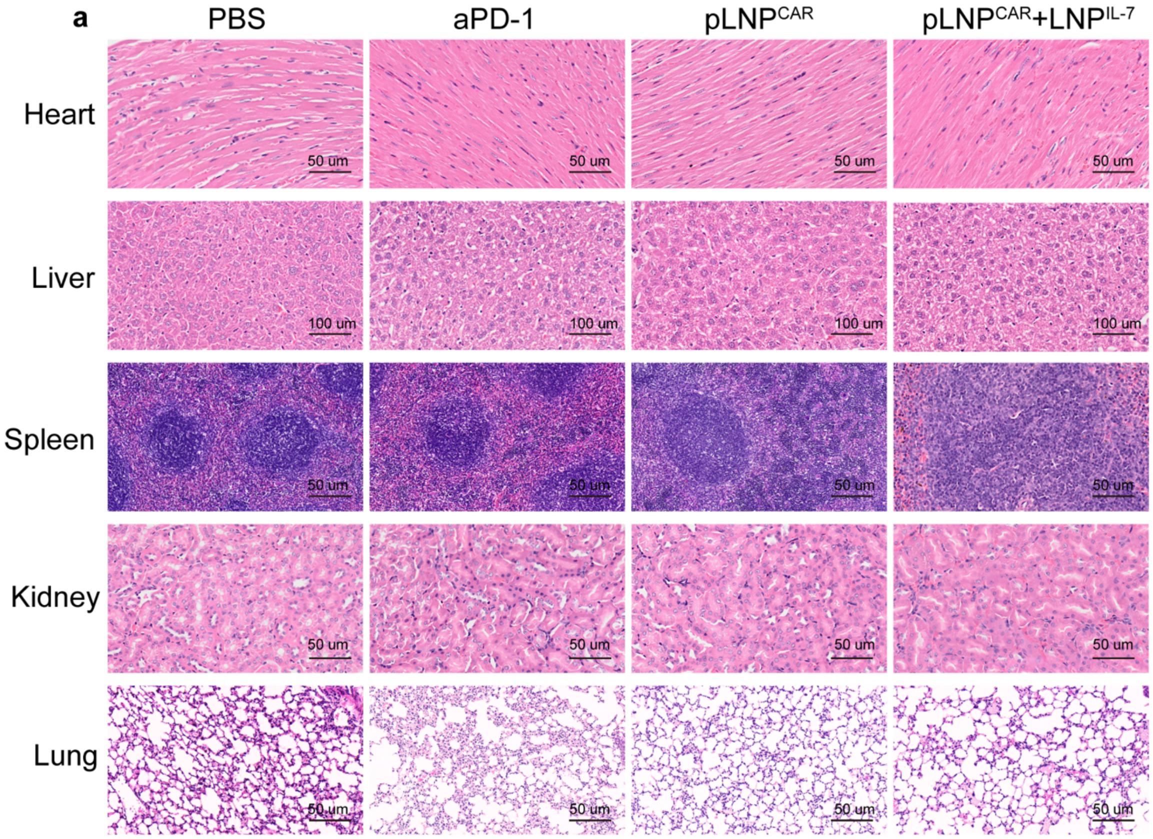


**Supplementary Fig. 5 Histology analysis of heart, liver, spleen, kidney and lung related to Fig. 5. a**, Representative H&E staining images from an experiment. Scale bar 100 µm for liver, 50 µm for the rest of the organs.

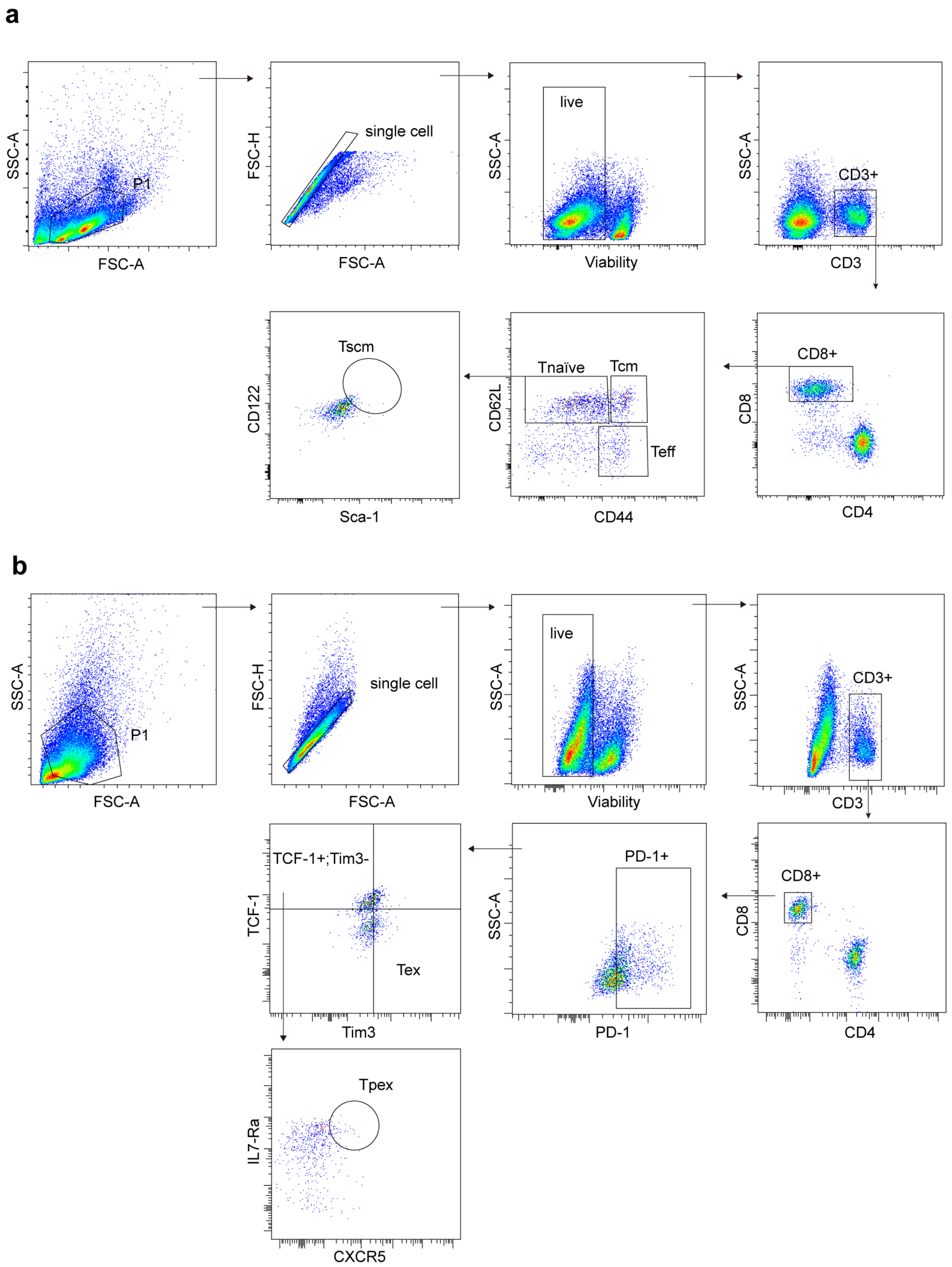


**Supplementary Fig. 6 Flow cytometry gating strategy related to Fig. 2 and Fig. 4.** Tnaïve, Tcm, Teff, Tscm cell gating strategy (**a**), Tpex and Tex gating strategy (**b**).
